# Flower initiates a positive feedback loop upon PIP_2_ enrichment at periactive zones to control bulk endocytosis

**DOI:** 10.1101/2020.06.22.164400

**Authors:** Tsai-Ning Li, Yu-Jung Chen, You-Tung Wang, Hsin-Chieh Lin, Chi-Kuang Yao

## Abstract

Synaptic vesicle (SV) endocytosis is coupled to exocytosis to maintain SV pool size and thus neurotransmitter release. Intense stimulation induces activity-dependent bulk endocytosis (ADBE) to recapture large quantities of SV constituents in large endosomes from which SVs reform. How these consecutive processes are spatiotemporally coordinated remains unknown. Here, we show that the Flower Ca^2+^ channel-dependent phosphatidylinositol 4,5-bisphosphate (PIP_2_) compartmentalization governs such control. Strong stimuli trigger PIP_2_ microdomain formation at periactive zones. Upon exocytosis Flower translocates from SVs to periactive zones, where it increases PIP_2_ levels via Ca^2+^ influxes. Remarkably, PIP_2_ directly enhances Flower channel activity, thereby establishing a positive feedback loop for PIP_2_ microdomain compartmentalization. The PIP_2_ microdomains drive ADBE and SV reformation from bulk endosomes. PIP_2_ further sorts Flower to bulk endosomes, thereby terminating endocytosis. Hence, we propose that the interplay between Flower and PIP_2_ is the crucial spatiotemporal cue that couples exocytosis to ADBE and subsequent SV reformation.

## Introduction

Proper synaptic vesicle (SV) exocytosis dictates the robustness of brain activity. Coupling SV exocytosis with proper endocytosis is crucial for maintaining a balance of SV proteins at the release site, plasma membrane equilibrium, SV identity, and SV pool size (Chanaday et al., 2019; Haucke et al., 2011; Lou, 2018; Wu et al., 2014a). Currently, four modes of SV endocytosis are proposed. These differ in terms of stimulation intensity for their induction, formation, and molecular components (Haucke et al., 2011; Kononenko and Haucke, 2015; Wu et al., 2014a). Under mild neuronal stimulation, the SV partially fuses with the plasma membrane and reforms at the active zone, the so called “kiss and run” mode. During clathrin-mediated endocytosis (CME), the SV fully collapses into the plasma membrane, followed by reformation of a single SV. Ultrafast endocytosis was also shown to recycle SVs at a sub-second timescale by forming a ~80 nm-sized bulk endosome predominantly at the edge of the active zone. SVs subsequently regenerate from this bulk endosome (Granseth et al., 2006; Watanabe et al., 2013a; Watanabe et al., 2013b; Zhu et al., 2009). High frequency stimulation and thus exocytosis could easily surpass the capacity of the three above-described endocytic mechanisms. It has therefore been proposed that activity-dependent bulk endocytosis (ADBE) has the necessary recapture capacity upon intense stimulation (Clayton et al., 2008; Soykan et al., 2017; Wu and Wu, 2007). This retrieval mode is elicited at the periactive zone to recapture large quantities of SV constituents via bulk endosome (~100-500 nm) formation, from which SVs subsequently reform. Hence, specific routes of SV recycling may fit the specific demands of a wide range of neuronal activities at the synapse. However, how stimulation intensity dictates the choice between these different endocytic modes is not well understood.

ADBE has been documented in many different types of neurons in invertebrates and vertebrates (Clayton et al., 2008; Heerssen et al., 2008; Heuser and Reese, 1973; Holt et al., 2003; Kasprowicz et al., 2008; Kittelmann et al., 2013; Miller and Heuser, 1984; Richards et al., 2000; Soykan et al., 2017; Stevens et al., 2012; Vijayakrishnan et al., 2009; Wenzel et al., 2012; Wu and Wu, 2007; Yao et al., 2017). Although the precise mechanism regulating ADBE is not known, multiple lines of evidence suggest that actin polymerization may serve as the membrane invagination force responsible for generating bulk endosomes (Gormal et al., 2015; Holt et al., 2003; Kokotos and Low, 2015; Nguyen et al., 2012; Richards et al., 2004; Soykan et al., 2017; Wu et al., 2016). Furthermore, phosphatidylinositol metabolism has also been implicated in controlling this recycling mode (Gaffield et al., 2011; Holt et al., 2003; Richards et al., 2004; Vijayakrishnan et al., 2009). Importantly, loss of Synaptojanin, the major phosphatidylinositol 4,5-bisphosphate (PIP_2_) catalytic enzyme in neurons (Tsujishita et al., 2001), leads to reduced SV endocytosis elicited by intense stimulation, presumably by affecting ADBE (Mani et al., 2007). PIP_2_ is known to promote actin polymerization by activating a number of actin modulators (Janmey et al., 2018). It has also been reported that formation of PIP_2_ microdomains precedes actin polymerization during a process reminiscent of ADBE in neurosecretory cells (Gormal et al., 2015). Thus, subcellular compartmentalization of PIP_2_ may provide the spatial information dictating where bulk membranes will invaginate. However, the mechanism initiating the formation of the PIP_2_ microdomains is unknown.

Coordinated SV protein and membrane retrieval plays an important role in maintaining the identity of newly formed SVs during SV recycling (Kaempf and Maritzen, 2017; McMahon and Boucrot, 2011; Saheki and De Camilli, 2012; Traub and Bonifacino, 2013). It has been well documented that proper sorting of SV proteins to the nascent SV is achieved in CME by the cooperative action of PIP_2_ and adaptor protein complexes (Saheki and De Camilli, 2012; Traub and Bonifacino, 2013). Recent studies have also revealed a distinct sorting mechanism that retrieves selective SV cargoes to the bulk endosome via ADBE (Kokotos et al., 2018; Nicholson-Fish et al., 2015). Several lines of evidence further suggest that clathrin and adaptor protein complexes are required for reforming SVs from the bulk endosome (Cheung and Cousin, 2012; Glyvuk et al., 2010; Kokotos et al., 2018; Kononenko et al., 2014; Park et al., 2016). A dynaminI/dynaminIII/clathrin-independent mechanism has also been reported as being involved in this process (Wu et al., 2014c). Thus, during SV regeneration via ADBE, multiple protein sorting steps may be required to ensure that the SVs harbor the proper compositions of lipids and proteins, thereby endowing specific release probabilities in relation to other modes of endocytosis (Cheung et al., 2010; Hoopmann et al., 2010; Nicholson-Fish et al., 2015; Silm et al., 2019). The mechanism by which protein sorting and membrane retrieval are coordinated in this process remains to be explored.

Here, we find that upon intense stimulation PIP_2_ is compartmentalized into microdomains at periactive zones in the synaptic boutons of *Drosophila* larval neuromuscular junctions (NMJs). Blockade of PIP_2_ microdomain formation diminishes ADBE and SV reformation from the bulk endosome. Increased intracellular Ca^2+^ and SV exocytosis are prerequisites for initiating ADBE (Morton et al., 2015; Wu and Wu, 2007). We have previously shown that Flower (Fwe), a SV-associated Ca^2+^ channel, regulates both CME and ADBE, and that its channel activity is strongly activated upon intense stimulation to elicit ADBE (Yao et al., 2017). We show that Fwe initiates a positive feedback koop upon PIP_2_ increase to ensure the formation of PIP_2_ microdomains and thus trigger ADBE and subsequent SV reformation. Intriguingly, Fwe is also selectively sorted to the bulk endosome by PIP_2_, thereby stopping membrane retrieval. Hence, spatiotemporal interplays between Flower and PIP_2_ coordinates retrieval of SV cargos and membranes, herein coupling exocytosis to ADBE and subsequent SV reformation.

## Results

### Intense neuronal activity induces formation of PIP_2_ microdomains at the presynaptic periactive zone of *Drosophila* synapses

To investigate the dynamics of PIP_2_ in the presynaptic compartment, we expressed a a GFP fusion protein of the pleckstrin homology (PH) domain of PLC_δ1_ (PLC_δ1_-PH-EGFP) in synaptic boutons of *Drosophila* larval NMJs using *nSyb-GAL4*, a pan-neuronal driver. PLC_δ1_-PH-EGFP binds to PIP_2_ with high affinity and is widely used to label subcellular compartments in which PIP_2_ is enriched (Chen et al., 2014; Khuong et al., 2010). We delivered 20-Hz stimuli for a few minutes to synaptic boutons in a 2-mM extracellular Ca^2+^ solution and performed live imaging. As shown in Figure 1a-b, we observed a very subtle increase in PLC_δ1_-PH-EGFP fluorescence in individual boutons (white arrows), similar to previous findings (Verstreken et al., 2009). However, when we raised the stimulus intensity to 40 Hz, we recorded a robust increase in fluorescence relative to a GFP fusion protein of the plasma membrane-integrated mCD8 domain (*UAS-mCD8-GFP*). Fluorescence signals rapidly returned to basal levels within tens of seconds when the stimuli were removed. We have previously documented that treatment with 40-Hz electric pulses or 90-mM high KCl solution can cause comparable stimulation intensities in *Drosophila* NMJ boutons (Yao et al., 2017). High K^+^ treatment also increased the fluorescence signal. No increase in the presynaptic protein level of PLC_δ1_-PH-EGFP was found under this condition (Figure 1-figure supplement 1a-b), arguing that this stimulation does not induce protein synthesis. These results suggest that, in response to intense stimulation, PLC_δ1_-PH-EGFP is localized and concentrated to PIP_2_-enriched microdomains, thereby enhancing the overall fluorescence.

**Figure 1.**
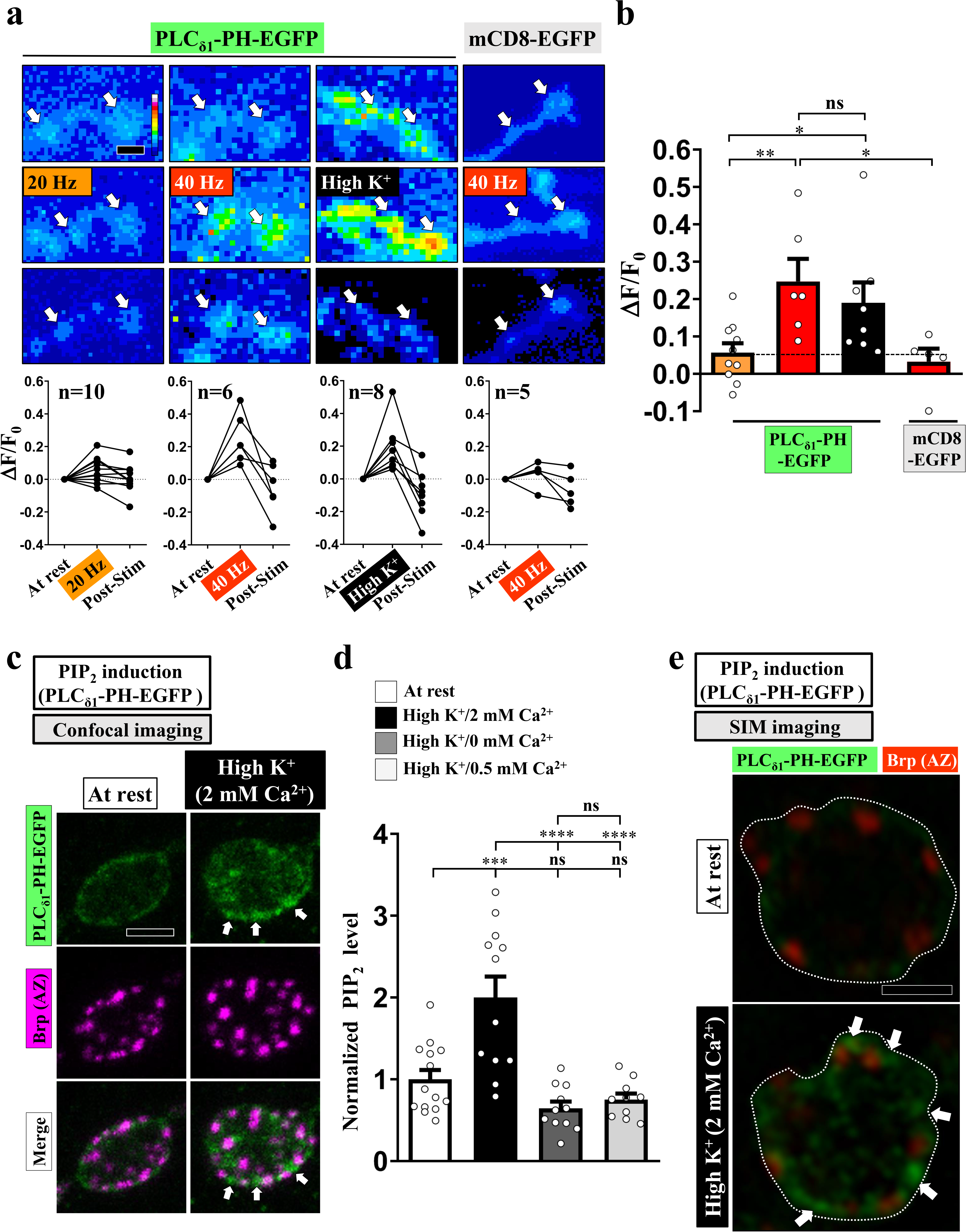
PIP_2_ forms microdomains at periactive zones under conditions of intense stimulation. (a-b) Increased fluorescence of PLC_δ1_-PH-EGFP but not mCD8-GFP in NMJ boutons upon intense stimulation. (a) (Top) Live images of the boutons (arrows) expressing *UAS-PLC_δ1_-PH-EGFP* or *UAS-mCD8-GFP*. Electrical (20 or 40 Hz) or chemical (90 mM K^+^) stimulation was conducted in a 2 mM-Ca^2+^ solution for 3 min (electrical) or 5 min (chemical) and then rested in 0 mM Ca^2+^ and 5 mM K^+^. Snapshot images taken before stimulation, at the third (electrical) and fifth (chemical) min of stimulation, and after stimulation. (Bottom) Traces of probe flourescence for single boutons. The number of boutons imaged (N). (b) Quantification data for EGFP fluorescence change. The resting fluorescence level (F_0_). Fluorescence change evoked by stimulation (ΔF). (c-e) PLC_δ1_-PH-EGFP is enriched at periactive zones dependently of Ca^2+^ upon intense stimulation. Confocal (c) or SIM (e) images of the boutons expressing PLC_δ1_-PH-EGF. The boutons subjected to high K^+^/2 mM Ca^2+^ (10-min stimulation of 90 mM K^+^/2 mM Ca^2+^), high K^+^/0 mM Ca^2+^ (10-min stimulation of 90 mM K^+^/0 mM Ca^2+^), or high K^+^/0.5 mM Ca^2+^ (1-min stimulation of 90 mM K^+^/0.5 mM Ca^2+^) treatments were fixed immediately and immunostained for PLC_δ1_-PH-EGFP (green) and Bruchpilot (Brp) [an active zone scaffold protein; magenta (c), red (e)]. The PLC_δ1_-PH-EGFP-enriched puncta (Arrows). (d) Quantification data for PLC_δ1_-PH-EGFP staining intensities, normalized to the value of the resting condition. Individual data values are shown in graphs. *P* values: ns, not significant; *, *P*<0.05; **, *P*<0.01; ***, *P*<0.001; ****, *P*<0.0001. Mean ± SEM. Scale bar: 1 μm (e), 2 μm (a, c). Statistics: one-way ANOVA with Tukey’s post hoc test (b, d). **Figure 1-source data 1. Source data for Figure 1**.

To characterize the subcellular distribution of the induced PIP_2_ microdomains, we developed a chemical fixation protocol whereby the NMJ boutons were fixed immediately after high K^+^ stimulation and immunostained for PLC_δ1_-PH-EGFP using an α-GFP antibody to enhance the signal. Confocal imaging revealed that, for neurons at rest, the majority of PLC_δ1_-PH-EGFP signal was dispersed throughout the cytosol, with some weak PLC_δ1_-PH-EGFP signals being associated with the presynaptic plasma membrane (Figure 1c). Upon stimulating with 90 mM K^+^ and 2 mM Ca^2+^ for 10 min, we observed high-level PLC_δ1_-PH-EGFP puncta close to the plasma membrane (Figure 1c-d) whereas the mCD8-GFP remained constant, both at rest and under conditions of high K^+^ stimulation (Figure 1-figure supplement 1c-d). The data indicate that PIP_2_ clusters in microdomains. Intriguingly, we noted that the induced PIP_2_ microdomains were primarily at periactive zones (Figure 1c), a hot-spot for ADBE (Chanaday et al., 2019; Kononenko and Haucke, 2015; Wu et al., 2014a). By using structured illumination microscopy (SIM), we estimate that the average size of these PIP_2_ microdomains is ~ 200 nm in diameter (Figure 1e)(0.0414 ± 0.004 μm^2^, mean ± S.E.M., n= 34 boutons).

Next, to determine the Ca^2+^ dependence of the PIP_2_ microdomains, we stimulated the boutons in a solution of 90 mM K^+^ and 0 mM Ca^2+^, which resulted in failure to induce PIP_2_ microdomain formation (Figure 1d). We obtained a similar result using 1-min stimulation of 90 mM K^+^ and 0.5 mM Ca^2+^ (Figure 1d), which was previously shown to primarily elicit CME but not ADBE (Yao et al., 2017). Hence, these results suggest that intense stimulation can elicit Ca^2+^-driven compartmentalization of PIP_2_ at the periactive zone in synaptic boutons of *Drosophila* NMJs.

### PIP_2_ microdomains are involved in ADBE initiation and SV reformation from bulk endosomes

Next, we investigated the function of PIP_2_ microdomains in ADBE. High K^+^ treatment is widely used to trigger ADBE in a broad range of synapses (Akbergenova and Bykhovskaia, 2009; Clayton et al., 2008; Jin et al., 2019a; Stevens et al., 2012; Vijayakrishnan et al., 2009; Wu and Wu, 2007; Wu et al., 2014c). To measure ADBE, we induced ADBE with 90 mM K^+^ and 2 mM Ca^2+^ for 10 min followed by transmission electron microscopy (TEM). ADBE was evoked in wild-type control boutons under these conditions and generated the formation of bulk endosomes (red asterisks, defined as > 80 nm in diameter) (Figure 2a, 2d). To suppress the function of PIP_2_, we expressed PLC_δ1_-PH-EGFP, anticipating that the PLC_δ1_-PH domain binding to PIP_2_ would restrict availability of PIP_2_ to its effectors and metabolic enzymes (Figure 2b) (Khuong et al., 2013). By using the *GAL4/UAS* system, we were able to adjust expression levels of the PLC_δ1_-PH domain by manipulating the temperature (Brand and Perrimon, 1993; D’Avino and Thummel, 1999; Wilder, 2000). When we neuronally express PLC_δ1_-PH-EGFP using *nSyb-GAL4* and grow larvae at 25 ºC, we found that mild expression of PLC_δ1_-PH-EGFP had a slightly inhibitory effect on ADBE induction relative to wild-type control boutons (Figure 2a, 2d). In contrast, when larvae were grown at 29 ºC, high PLC_δ1_-PH-EGFP expression almost completely abolished ADBE (Figure 2a, 2d).

**Figure 2.**
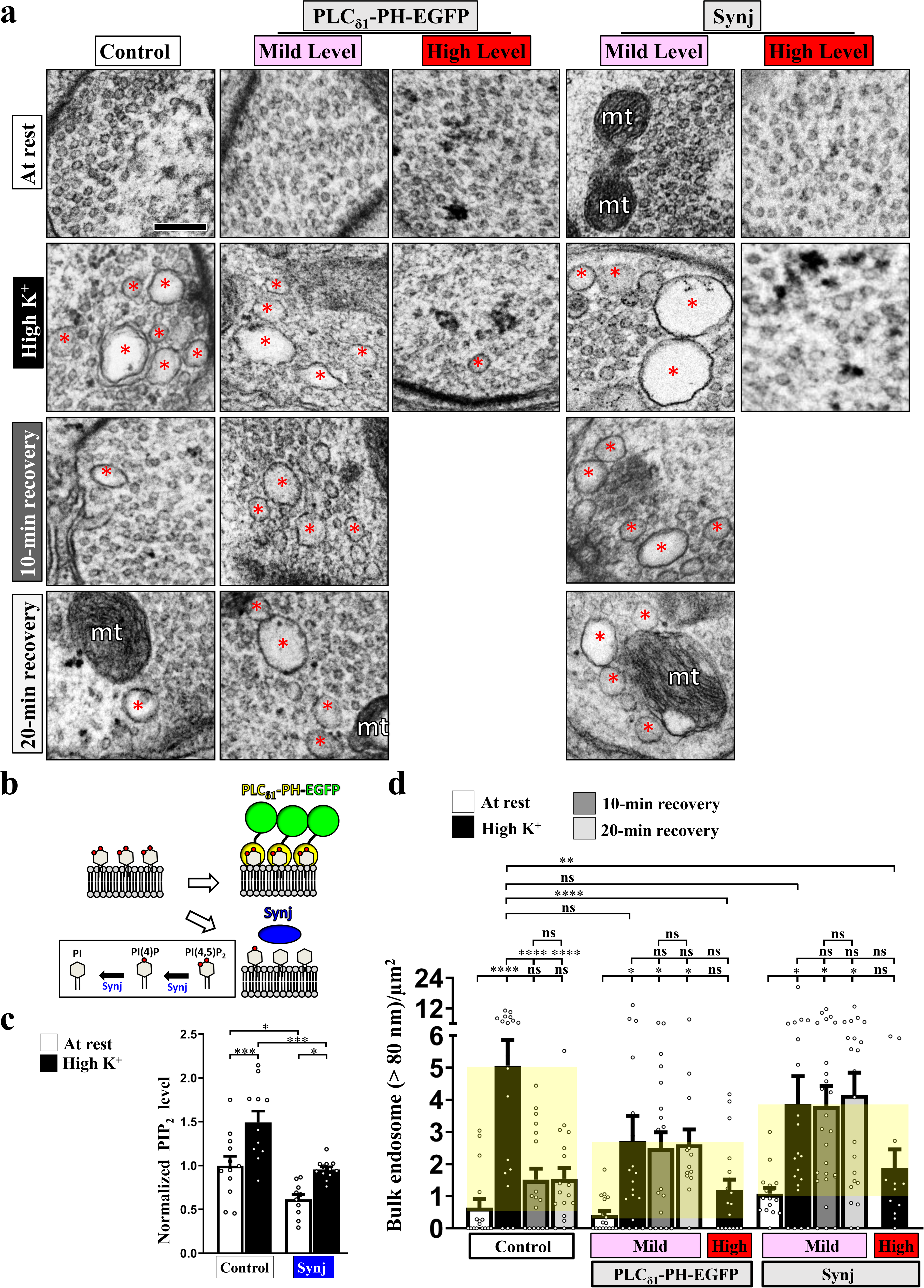
PIP_2_ microdomains drive ADBE and SV reformation from bulk endosomes. Reducing PIP_2_ availability suppresses ADBE and subsequent SV reformation. (a) TEM images of the boutons of controls (*nSyb-GAL4/+*), mild PLC_δ1_-PH-EGFP expression (*nSyb-GAL4/UAS-PLC_δ1_-PH-EGFP* at 25 ºC), high PLC_δ1_-PH-EGFP expression (*nSyb-GAL4/UAS-PLC_δ1_-PH-EGFP* at 29 ºC), mild Synj expression (*nSyb-GAL4/UAS-synj* at 25 ºC), or high Synj expression (*nSyb-GAL4/UAS-synj* at 29 ºC). At rest (10-min incubation of 5 mM K^+^/0 mM Ca^2+^). High K^+^ (10-min stimulation of 90 mM K^+^/2 mM Ca^2+^). 10-min recovery (10-min stimulation of 90 mM K^+^/2 mM Ca^2+^, followed by 10-min incubation of 5 mM K^+^/0 mM Ca^2+^). 20-min recovery (10-min stimulation of 90 mM K^+^/2 mM Ca^2+^, followed by 20-min incubation of 5 mM K^+^/0 mM Ca^2+^). Bulk endosomes (> 80 nm in diameter, red asterisks). Mitochondria (mt). Quantification data for total number of bulk endosomes per bouton area (d). (b) A schematic for PIP_2_ suppression by PLC_δ1_-PH-EGFP and Synj expression. (c) Expression of Synj reduced presynaptic PIP_2_ at rest and in high K^+^. The boutons co-expressing *UAS-PLC_δ1_-PH-EGFP* with *UAS-RFP* (control) or *UAS-synj* using *nSyb-GAL4* were subjected to resting condition (10-min incubation of 5 mM K^+^/0 mM Ca^2+^) or high K^+^ stimulation (10-min stimulation of 90 mM K^+^/2 mM Ca^2+^), followed by α-GFP immunostaining. Confocal images of the boutons (Figure 2-figure supplement 1a). Quantification data for PLC_δ1_-PH-EGFP staining intensity are shown, normalized to the value of the resting condition of controls. Individual data values are shown in graphs. *P* values: ns, not significant; *, *P*<0.05; **, *P*<0.01; ***, *P*<0.001; ****, *P*<0.0001. Mean ± SEM. Scale bar: 500 nm. Statistics: one-way ANOVA with Tukey’s post hoc test. **Figure 2-source data 1. Source data for Figure 2**.

Synaptojanin (Synj) functions as the major neuronal PIP_2_ phosphatase in mammals (Tsujishita et al., 2001) and *Drosophila* (Chen et al., 2014; Verstreken et al., 2009). Synj comprises a central 5-phosphatase domain that specifically dephosphorylates the 5’ position of PI(4,5)P_2_ to produce PI(4)P (McPherson et al., 1996; Woscholski et al., 1997). In addition, an N-terminal Sac1 domain converts several phosphatidylinositides—including PI(3,5)P_2_, PI(3)P, and PI(4)P—to PI (Figure 2b) (Guo et al., 1999). Indeed, overexpression of Synj lowers PIP_2_ levels at synaptic boutons at resting state, and diminishes the formation of the PIP_2_ microdomains induced by high K^+^ (Figure 2c; Figure 2-figure supplement 1a). Thus, overexpression of Synj exerts a dosage-dependent inhibition of ADBE (Figure 2a, 2d), similar to the effect of the increase in PLC_δ1_-PH. To assess if this ADBE deficiency may arise, in part, from insufficient SV exocytosis (Morton et al., 2015), we performed a styryl FM dye loading and unloading assay to measure exocytosis. SV exocytosis proceeded normally in boutons expressing PLC_δ1_-PH-EGFP or Synj relative to *nSyb-GAL4* controls (Figure 2-figure supplement 2). Hence, the PIP_2_ microdomains play an essential role in inducing ADBE.

SVs regenerate from bulk endosomes within minutes of their formation (Cheung and Cousin, 2012; Glyvuk et al., 2010; Kononenko et al., 2014; Stevens et al., 2012; Wu et al., 2014c). Using an approach employed previously (Stevens et al., 2012), we treated the boutons with high K^+^ followed by an incubation in 5 mM K^+^ and 0 mM Ca^2+^ solution for 10 or 20 minutes, allowing the SVs to reform from the bulk endosomes. In controls (*nSyb-GAL4)*, both the number and area of the induced bulk endosomes reverted to almost basal levels within 10 min (Figure 2a, 2d; Figure 2-figure supplement 1b), demonstrating proper SV reformation. Interestingly, upon expression of low levels of PLC_δ1_-PH-EGFP, the induced bulk endosomes remained after 10 min and 20 min recovery periods, yet ADBE was only partially impaired (Figure 2a, 2d; Figure 2-figure supplement 1b). A strong retention of SV reformation ability was also found upon mild overexpression of Synj (Figure 2a, 2d; Figure 2-figure supplement 1b). Hence, the PIP_2_ microdomains differentially control ADBE and SV reformation from the bulk endosome, with the latter event being elicited by relatively high levels of PIP_2_.

### PIP_2_ microdomains are established via a positive feedback loop of Fwe and PIP_2_

We have previously shown that the SV-associated Ca^2+^ channel Flower (Fwe) elevates presynaptic Ca^2+^ levels in response to strong stimuli to trigger ADBE (Yao et al., 2009; Yao et al., 2017). Given the Ca^2+^ dependence of PIP_2_ microdomains (Figure 1d), we hypothesized that exocytosis evoked by intense stimulation promotes Fwe clustering at periactive zone, where it may provide the Ca^2+^ influx to induce PIP_2_ microdomain formation. To test this hypothesis, we first conducted a proximity ligation assay (PLA) (Soderberg et al., 2008) to investigate if there was a close association between Fwe and PIP_2_ in response to stimulation. In our PLA (Figure 3a), *UAS-Flag-Fwe-HA* was expressed in a *fwe* mutant background to replace endogenous Fwe protein with a tagged protein, and expression of *UAS-PLC_δ1_-PH-EGFP* reported the localization of PIP_2_. Primary antibodies against the HA tag and GFP protein were used to detect interactions between Flag-Fwe-HA and PLC_δ1_-PH-EGFP. The PLA signal was low in resting boutons. In contrast, high K^+^ treatment significantly increased PLA signal intensity (Figure 3a-b) suggesting a physical interaction between Fwe and PIP_2_.

**Figure 3.**
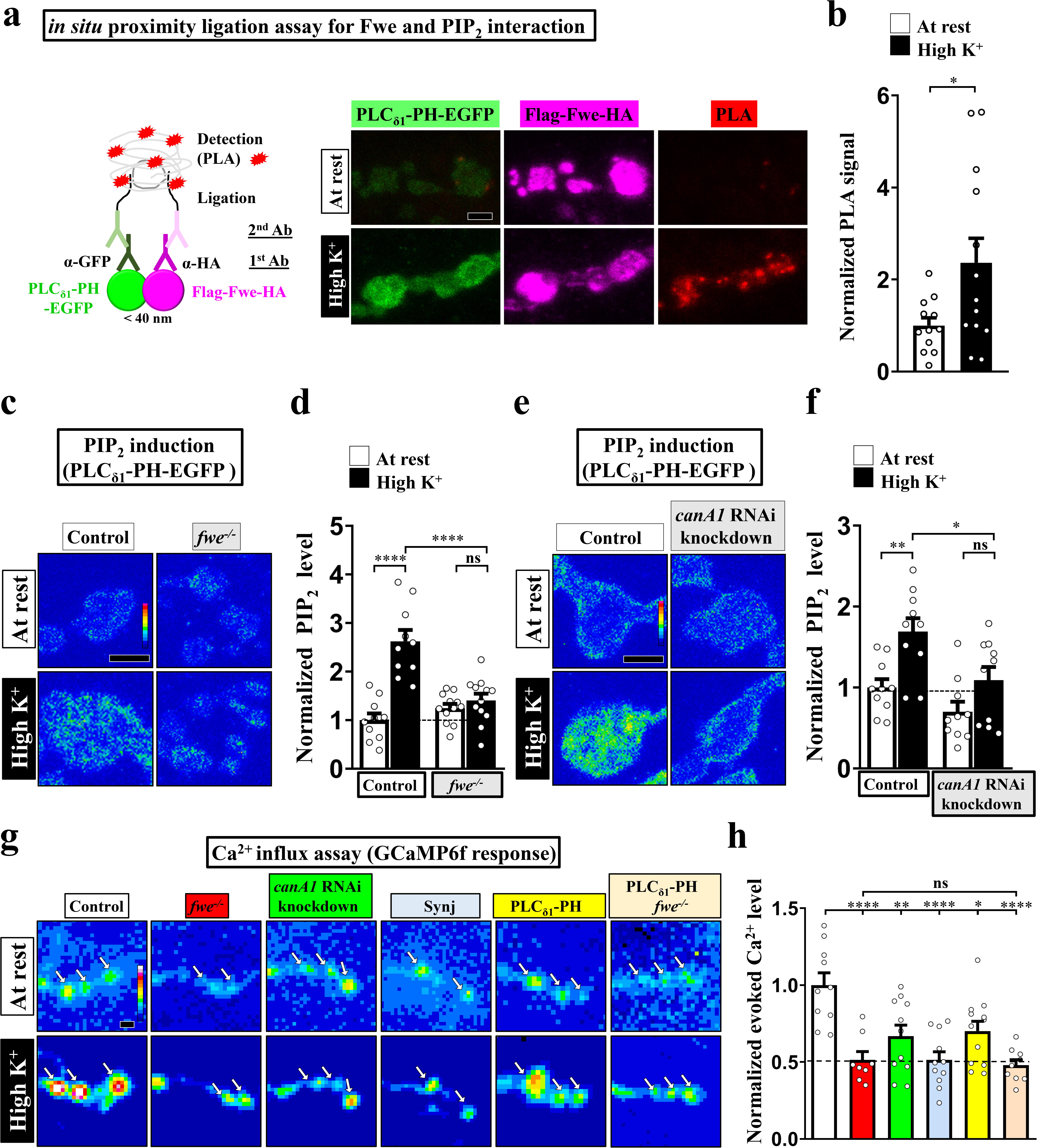
Fwe and PIP_2_ form a positive feedback loop to establish PIP_2_ microdomains. (a-b) Fwe and PIP_2_ interact upon intense stimulation. (a) (Left) A schematic for PLA. (Right) Confocal images of *fwe* mutant boutons co-expressing Flag-Fwe-HA and PLC_δ1_-PH-EGFP (*nSyb-GAL4/UAS-Flag-fwe-HA*/ *UAS-PLC_δ1_-PH-EGFP in fwe^DB25/DB56^*). After resting condition or high K^+^ stimulation, the boutons were subjected to PLA. α-GFP and α-HA stained for PLC_δ1_-PH-EGFP (green) and Flag-Fwe-HA (magenta), respectively. PLA signals (red). (b) Quantification data for PLA signal intensities, normalized to the value of resting condition. (c-f) *fwe* mutation or *canA1* RNAi knockdown perturbs PIP_2_ microdomain formation. (c) Pseudocolored confocal images of the boutons neuronally expressing *UAS-PLC_δ1_-PH-EGFP* in *fwe^DB25/+^* or *fwe^DB25/DB56^*. (e) Pseudocolored confocal images of the boutons co-expressing *UAS-PLC_δ1_-PH-EGFP* with *UAS-RFP* or *UAS-canA1FB5* using *nSyb-GAL4*. After resting condition or high K^+^ stimulation, the boutons were stained with α-GFP. (d and f) Quantification data for PLC_δ1_-PH-EGFP staining intensities, normalized to the value of the resting condition of controls. (g-h) Impaired PIP_2_ microdomain attenuates Fwe Ca^2+^ conductance. (g) Snapshot Ca^2+^ images of the boutons (arrows) expressing *lexAOP2-GCaMP6f* using *vglut-lexA*. Control (*w^1118^*), *fwe* mutant (*fwe^DB25/DB56^*). *canA1* knockdown (*nSyb-GAL4/UAS-canA1FB5*). Synj overexpression (*nSyb-GAL4/UAS-synj*). PLC_δ1_-PH-APEX2-HA expression (*vglut-lexA/LexAOP2-PLC_δ1_-PH-APEX2-HA*). *fwe* mutant expressing PLC_δ1_-PH-APEX2-HA (*vglut-lexA/LexAOP2-PLC_δ1_-PH-APEX2-HA* in *fwe^DB25/DB56^*). Imaging was taken in the fifth minute after high K^+^ (2 mM Ca^2+^) stimulation. (h) Quantification data for evoked Ca^2+^ level, normalized to the value of controls. Evoked Ca^2+^ levels are shown as the increase in GCaMP6f fluorescence under high K^+^ stimulation. Individual data values are shown in graphs. *P* values: ns, not significant; *, *P*<0.05; **, *P*<0.01; ****, *P*<0.0001. Mean ± SEM. Scale bar: 2 μm. Statistics: Student *t*-test (b). One-way ANOVA with Tukey’s post hoc test (d, f, h). **Figure 3-source data 1. Source data for Figure 3**.

Next, we investigated the effect of loss of Fwe on PIP_2_ microdomain formation. Basal levels of PIP_2_ were not affected in the *fwe* mutant relative to control (Figure 3c-d). However, intense stimulation failed to elicit PIP_2_ microdomain formation in the *fwe* mutant (Figure 3c-d), arguing that Fwe triggers the formation of PIP_2_ microdomains. Calmodulin and Calcineurin are thought to be the Ca^2+^ sensors for ADBE (Evans and Cousin, 2007; Jin et al., 2019b; Marks and McMahon, 1998; Sun et al., 2010; Wu et al., 2009; Wu et al., 2014b). The Calcineurin complex dephosphorylates PIP_2_ metabolic enzymes, including Synj and Phosphatidylinositol 4-phosphate 5-kinase Iγ (PIPKIγ) (Cousin and Robinson, 2001; Lee et al., 2005; Lee et al., 2004). *Drosophila* possesses three isoforms of the catalytic subunit of Calcineurin, i.e., CanA1, CanA-14F, and Pp2B-14D. CanA1 regulates development of *Drosophila* NMJ boutons (Wong et al., 2014). We therefore knocked down *canA1* in neurons by expressing *UAS-canA1FB5*, a specific *canA1-RNAi* construct (Dijkers and O’Farrell, 2007; Wong et al., 2014). Similar to the effect of loss of Fwe, reduced CanA1 levels greatly suppressed the formation of PIP_2_ microdomains compared to the control (Figure 3e-f), indicating that Calcineurin mediates the Fwe Ca^2+^ channel to induce PIP_2_ microdomains.

Our previous Ca^2+^ imaging data showed that intense stimulation activates Fwe to increase presynaptic Ca^2+^ concentrations (Yao et al., 2017). We measured intracellular Ca^2+^ levels with the Ca^2+^ indicator GCaMP6f (Chen et al., 2013), and found that whereas control boutons exhibited a robust increase in intracellular Ca^2+^ upon high K^+^ stimulation, *fwe* mutant boutons exhibit an impaired Ca^2+^ response (Figure 3g-h). Unexpectedly, *canA1* knockdown also elicited the same deficient Ca^2+^ response (Figure 3g-h). Similar suppressive effects were obtained upon overexpressing Synj or the PLC_δ1_-PH domain (Figure 3g-h). Hence, Calcineurin activation may increase PIP_2_ activity, which may in turn promote the Ca^2+^ channel activity of Fwe to further increase intracellular Ca^2+^ levels. To investigate the potential for such feedback regulation, we expressed the PLC_δ1_-PH domain in a loss of Fwe background. Expression of the PLC_δ1_-PH domain did not rescue the low Ca^2+^ concentration caused by the *fwe* mutation (Figure 3g-h), showing that the Ca^2+^ suppression exerted by the PLC_δ1_-PH domain indeed depends on Fwe. These findings argue that a positive feedback loop involving Fwe and PIP_2_ is responsible for the formation of PIP_2_ microdomains.

### PIP_2_ gates Fwe

A well-known function of PIP_2_ is to modulate ion channel activity through its electrostatic binding to clustered positively-charged amino acids adjacent to the transmembrane domains of ion channels (Hille et al., 2015; Suh and Hille, 2008). Through protein alignment analysis, we found tandem positively-charged amino acids, including lysine (K) and arginine (R), in the intracellular juxta-transmembrane regions of Fwe. These residues are evolutionarily conserved in mice and humans (Figure 4a), whereas other cytosolic residues show poor conservation. This feature inspired us to test the potential impact of PIP_2_ on the channel function of Fwe. To determine direct interaction between Fwe and PIP_2_, we conducted a nanoluciferase (Nluc)-based bioluminance resonance energy transfer (BRET) assay (Cabanos et al., 2017). In our BRET assay (Figure 4a), upon ion channel binding of BODIPY-TMR-conjugated PIP_2_, illumination of Nluc-fused ion channels wrapped in detergent-formed micelles can excite BODIPY-TMR-conjugated PIP_2_ to emit a BRET signal. As shown in Figure 4b, we reconstituted the micelles containing purified Nluc-Fwe-1D4 fusion proteins and, after adding BODIPY-TMR-PIP_2_ and furimazine (a Nluc substrate), we observed a remarkable increase in BRET signal. Excess cold PIP_2_ reduced the signal to ~50% of the BRET signals by competing for the PIP_2_ binding sites in Fwe, suggesting direct PIP_2_ binding to Fwe. To assess the involvement of the positively-charged amino acids of Fwe in PIP_2_ binding, we mutated the residues to non-charged alanine to eliminate the electrostatic interactions. Residue substitution resulted in a significant reduction in BRET signal, comparable to the competitive effect attributable to provision of excess cold PIP_2_. Therefore, the majority of the PIP_2_ binding activity of Fwe is mediated by these positively-charged amino acids. Furthermore, alanine substitution of residues in both the middle (K95/K100/R105) and C-terminal (K146/K147/R150) regions of Fwe reduced PIP_2_ specific binding (Figure 4b-c). Moreover, when N-terminal residues (K29/R33) of Fwe were further substituted with alanines, binding of PIP_2_ was also reduced (Figure 4b-c). Hence, these *in vitro* assays reveal that Fwe directly binds PIP_2_ through multiple regions.

**Figure 4.**
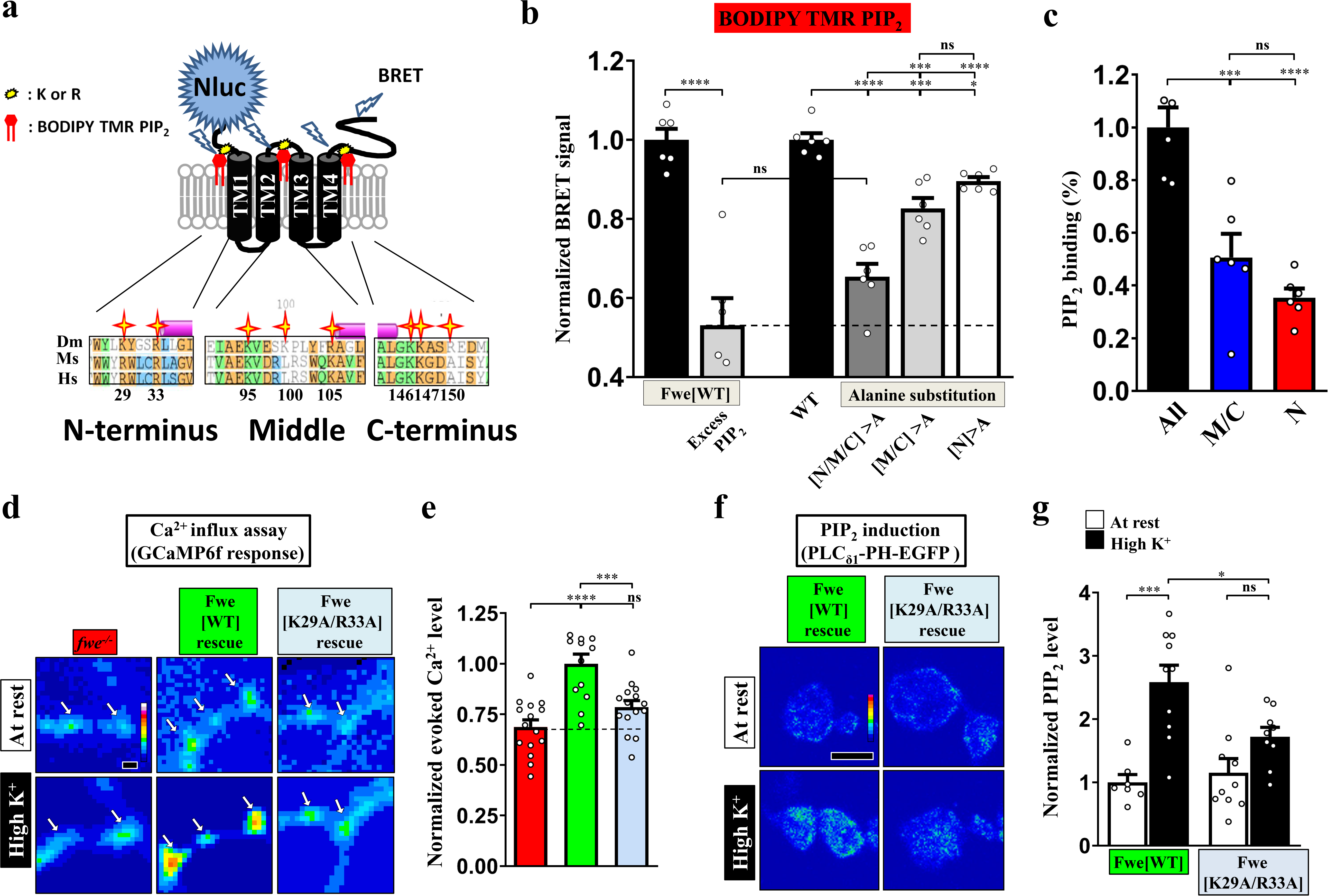
PIP_2_ binds to Fwe and promotes its Ca^2+^ channelling activity. (a-c) PIP_2_ binds Fwe. (a) A schematic of the Fwe structure and a BRET assay. Stars highlight conserved lysine (K) and arginine (R) in juxta-transmembrane regions: N-terminus (N), middle-domain (M), and C-terminus (C). *Drosophila Melanogaster* (Dm). *Mus Musculus* (Ms). *Human Sapiens* (Hs). N-terminal fusion of Nluc to Fwe allows to detect PIP_2_ binding. A 1D4 epitode is used for protein purification. (b) Nluc-Fwe-1D4 in micelles excites BODIPY-TMR-conjugated PIP_2_ to emit BRET signal, which is decreased by competitive cold PIP_2_ (1 mM). Alanine substitution of positively-charged residues in all regions (N/M/C>A), both middle-domain and C-terminus (M/C>A), or N-terminus (N>A) reduced BRET signals. (c) Corresponding PIP_2_ binding ability was calculated by subtracting the signal values of N/M/C>A, M/C>A or N>A from that for WT. Quantification data was normalized to the signal value of N/M/C. (d) PIP_2_ controls Fwe channel activity. Snapshot Ca^2+^ images of the boutons (arrows) expressing *lexAOP2-GCaMP6f* using *vglut-lexA*. *fwe* mutant (*fwe^DB25/DB56^*). HA-Fwe[WT]-APEX2 rescue (*nSyb-GAL4/UAS-HA-Fwe[WT]-APEX2 in fwe^DB25/DB56^*). HA-Fwe[K29/R33A]-APEX2 rescue (*nSyb-GAL4/UAS-HA-Fwe[K29/R33A]-APEX2 in fwe^DB25/DB56^*). Imaging was taken in the fifth minute after high K^+^ stimulation. (e) Quantification data for evoked Ca^2+^ level, normalized to the value of HA-Fwe-APEX2 rescue larvae. (f) Pseudocolored confocal images of the boutons expressing *UAS-PLC_δ1_-PH-EGFP*. After resting conditions or high K^+^ stimulation, the boutons were stained with α-GFP. (g) Quantification data for the PLC_δ1_-PH-EGFP staining intensity, normalized to the value of resting condition of HA-Fwe-APEX2 rescue larvae. Individual data values are shown in graphs. *P* values: ns, not significant; *, *P*<0.05; ***, *P*<0.001; ****, *P*<0.0001. Mean ± SEM. Scale bar: 2 μm (d, f). Statistics: Student *t*-test (b). One-way ANOVA with Tukey’s post hoc test (b, c, e, g). **Figure 4-source data 1. Source data for Figure 4**.

To directly test how PIP_2_ affects Flower Ca^2+^ channel function, we generated *UAS* transgenes for the Fwe variants in which all or subsets of positively-charged amino acids were mutated to alanine and performed a mutant rescue experiment using *nSyb-GAL4*. Mutations of all nine residues or only those in the middle region (K95/K100/R105) led to very lowly expressed protein levels, preventing further study. However, alanine substitution of C-terminal residues K146/K147/R150 did not affect SV localization of Fwe or its ability to regulate presynaptic Ca^2+^ concentration and induce PIP_2_ microdomain formation (Figure 4-figure supplement 1a-f), suggesting that these residues do not play a regulatory role in Fwe channel activity. We mutated the N-terminal residues K29/R33. The resulting K29A/R33A variant was still able to properly localize to presynaptic terminals (Figure 4-figure supplement 2a-b). However, upon high K^+^ stimulation, the K29A/R33A variant lost that ability to maintain proper intracellular Ca^2+^ levels (Figure 4d-e). Moreover, that variant failed to promote PIP_2_ microdomain formation upon high K^+^ stimulation (Figure 4f-g). These results reveal that the positive feedback loop involving Fwe and PIP_2_ relies on PIP_2_-dependent gating control of Fwe.

### Blockade of the positive feedback loop reduces ADBE and SV reformation from bulk endosomes

Next, we assessed the impact of the Fwe and PIP_2_ regulatory feedback loop on ADBE. Loss of Fwe severely impaired formation of bulk endosomes induced by ADBE under high K^+^ conditions compared to the *fwe* mutant rescue control (Figure 5a-b), consistent with our previous findings (Yao et al., 2017). Neuronal knockdown of *canA1* also blocked ADBE relative to the *nSyb-GAL4* control (Figure 5a-b). Expression of wild-type Fwe protein or the K146A/K147A/R150A mutant variant restored proper ADBE in the *fwe* mutant background (Figure 5a-b; Figure 4-figure supplement 1g-h), whereas expression of the K29A/R33A variant failed to rescue deficient ADBE (Figure 5a-b). Consistent with the suppressive effects caused by expression of the PLCδ1-PH domain or Synj (Figure 2), we found that all of the bulk endosomes that remained in boutons lacking *fwe* could not generate new SVs during a 10-min and even 20-min recovery period (Figure 5a-b; Figure 5-figure supplement 1). Expression of wild-type Fwe protein but not the K29A/R33A mutant variant rescued this defect (Figure 5a-b; Figure 5-figure supplement 1). Taken together with our results reported in previous sections, we propose that the positive feedback loop involving Fwe and PIP_2_ compartmentalizes PIP_2_ microdomains at the periactive zone of boutons to dictate and coordinate ADBE and subsequent SV reformation.

**Figure 5.**
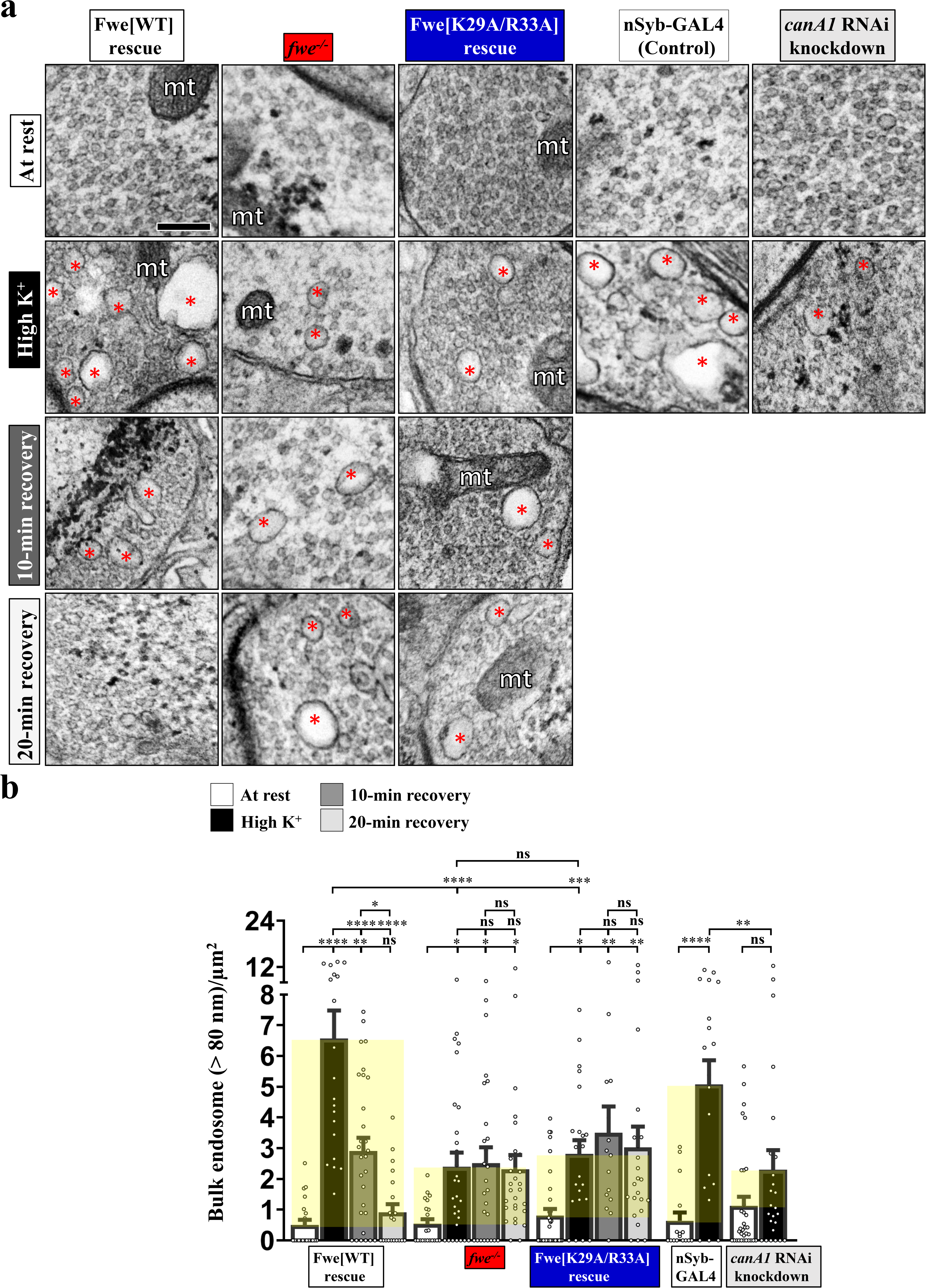
Perturbations of the positive feedback loop involving Fwe and PIP_2_ suppress ADBE and SV reformation from bulk endosomes. TEM images of the boutons of HA-Fwe-APEX2 rescue (*nSyb-GAL4/UAS-HA-Fwe-APEX2 in fwe^DB25/DB56^*), *fwe* mutant (*fwe^DB25/DB56^*), HA-Fwe[K29/R33A]-APEX2 rescue (*nSyb-GAL4/UAS-HA-Fwe[K29/R33A]-APEX2 in fwe^DB25/DB56^*), control (*nSyb-GAL4*), or *canA1* RNAi knockdown (*nSyb-GAL4/UAS-canA1FB5*). TEM processing was performed after the following treatments: at rest (10-min incubation of 5 mM K^+^/0 mM Ca^2+^); high K^+^ stimulation (10-min stimulation of 90 mM K^+^/2 mM Ca^2+^); 10-min recovery (10-min stimulation of 90 mM K^+^/2 mM Ca^2+^, followed by 10-min incubation of 5 mM K^+^/0 mM Ca^2+^); or 20-min recovery (10-min stimulation of 90 mM K^+^/2 mM Ca^2+^, followed by 20-min incubation of 5 mM K^+^/0 mM Ca^2+^). Bulk endosomes (>80 nm in diameter, red asterisks). Mitochondria (mt). (b) Quantification data of total numbers of bulk endosomes per bouton area. Individual data values are shown in graphs. *P* values: ns, not significant; *, *P*<0.05; **, *P*<0.01; ***, *P*<0.001; ****, *P*<0.0001. Mean ± SEM. Scale bar: 500 nm. Statistics: one-way ANOVA with Tukey’s post hoc test. **Figure 5-source data 1. Source data for Figure 5**.

### PIP_2_ facilitates sorting of Fwe to bulk endosomes

It was reported recently that a SV protein sorting process occurs during ADBE (Kokotos et al., 2018; Nicholson-Fish et al., 2015). VAMP4, a v-SNARE protein, is essential for ADBE to proceed, and it is selectively retrieved by ADBE (Kokotos et al., 2018; Nicholson-Fish et al., 2015). Given the important role of Fwe in triggering ADBE, we wondered if Fwe is sorted to bulk endosomes during ADBE. To visualize the vesicular localization of Fwe, we expressed a *UAS* transgene of the APEX2 fusion protein of Fwe (*UAS-HA-Fwe-APEX2*) in the *fwe* mutant background followed by diaminobenzidine (DAB) labelling and TEM. APEX2 is an engineering peroxidase that is capable of catalyzing DAB polymerization and proximal deposition, with the DAB polymers binding electron-dense osmium to enhance electron microscopy contrast (Lam et al., 2015). We found no specific DAB staining in Flag-Fwe-HA-rescued control boutons under high K^+^ stimulation conditions (Figure 6a). In HA-Fwe-APEX2-rescued boutons stimulated with high K^+^, the SV-based localization of Fwe was clearly revealed by DAB staining (white arrows, Figure 6b-c), consistent with our previous immunogold staining data (Yao et al., 2009). In addition, DAB signals are apparent on all bulk endosomes. Remarkably, the DAB staining intensities on a large portion (~ 75%) of bulk endosomes were much stronger than those on SVs (Figure 6b-c, 6f), suggesting a mechanism by which Fwe is predominantly sorted to bulk endosomes after Fwe initiates ADBE.

**Figure 6.**
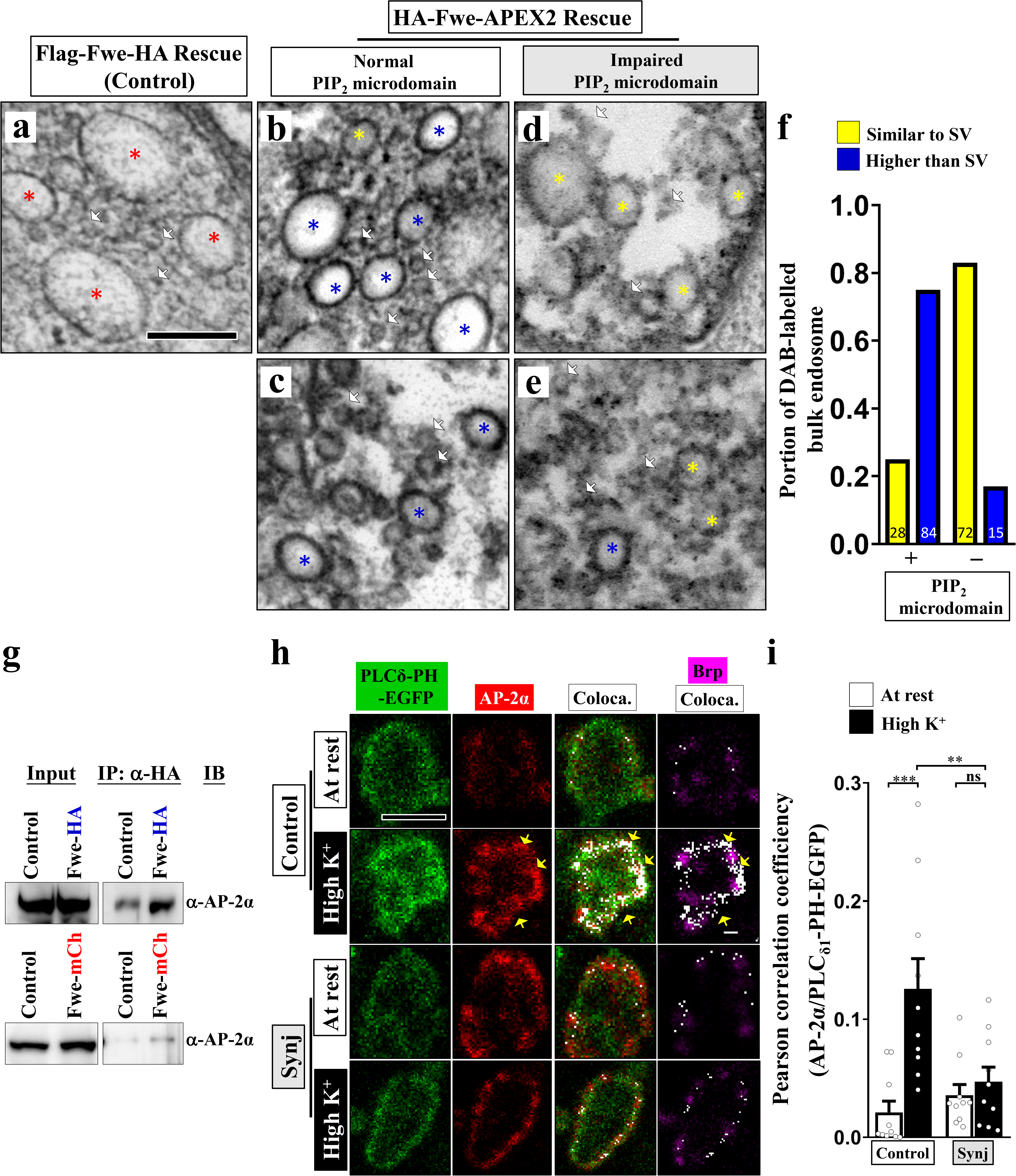
PIP_2_ and the AP-2 complex facilitate sorting of Fwe to bulk endosomes. (a-f) PIP_2_ sorts Fwe to bulk endosomes. (a-e) TEM images of the boutons of Flag-Fwe-HA rescue (*nSyb-GAL4/UAS-Flag-Fwe-HA in fwe^DB25/DB56^*), HA-Fwe-APEX2 rescue (*nSyb-GAL4/UAS-HA-Fwe-APEX2 in fwe^DB25/DB56^*), or HA-Fwe-APEX2 rescue coexpressing Synj (*nSyb-GAL4/UAS-HA-Fwe-APEX2*/ *UAS-synj in fwe^DB25/DB56^*). After high K^+^ stimulation, the boutons were subjected to DAB labeling. SVs (white arrows). DAB-negative bulk endosomes (red asterisks). Bulk endosomes with similar DAB levels to surrounding SVs (yellow asterisks) with higher DAB levels than those of SVs (blue asterisks). Perturbing PIP_2_ micordomain formation by Synj expression reduced DAB labeling of bulk endosomes. (f) Quantification data for DAB-labelled bulk endosomes. The number of bulk endosomes counted from ≥ 13 boutons is shown in graphs. (g) Fwe interacts with the AP-2 complex *in vivo*. Inputs account for 2% of pulldown. (h and i) PIP_2_ recruits AP-2 complexes. (h) Confocal images of the boutons co-expressing *UAS-PLC_δ1_-PH-EGFP* with *UAS-lacZ* or *UAS-synj* using *nSyb-GAL4*. After resting condition or high K^+^ stimulation, the boutons were stained for PLC_δ1_-PH-EGFP (green), AP-2α (red), and Brp (magenta). White areas represent overlapping regions of PLC_δ1_-PH-EGFP and AP-2α signals, which was significantly enhanced at the periactive zone upon high K^+^ stimulation. Synj overexpression impaired recruitment of the AP-2 complex. (i) Quantification data for Pearson correlation coefficients, calculated by normalizing the overlapping area of PLC_δ1_-PH-EGFP and AP-2α to the area of individual boutons. Individual data values are shown in graphs. *P* values: ns, not significant; **, *P*<0.01; ***, *P*<0.001. Mean ± SEM. Scale bar: 500 nm (a-e), 2 μm (h). Statistics: one-way ANOVA with Tukey’s post hoc test. **Figure 6-source data 1. Source data for Figure 6**.

PIP_2_ is known to recruit adaptor protein complexes to sort SV proteins to the nascent SV during CME (Saheki and De Camilli, 2012). To investigate if a similar mechanism participates in retrieval of Fwe during ADBE, we introduced a HA-tagged or mCherry-tagged Fwe genomic rescue transgene into an *fwe* mutant background and conducted co-immunoprecipitation experiments. Immunoprecipitation of the HA- or mCherry-tagged Fwe protein from adult head extracts pulled down the α subunit of the AP-2 complex (AP-2α) (Figure 6g), arguing that Fwe interacts with AP-2 complex, in addition to binding PIP_2_. The AP-2 adaptor complex is known to act in tandem with PIP_2_ (McMahon and Boucrot, 2011). Next, we examined the spatial relationship between PIP_2_ microdomains and AP-2 complex. In control boutons at rest, immunostaining of AP-2α revealed that AP-2 complexes were dispersed in the presynaptic compartment. However, upon high K^+^ stimulation, AP-2 complexes were localized to the periactive zone, where they were strongly associated with PIP_2_ microdomains (Figure 6h-i). Notably, Synj overexpression perturbed the localization of AP-2 complex at the periactive zone and PIP_2_ microdomains (Figure 6h-i), revealing that PIP_2_ enrichment in microdomains results in recruitment of AP-2 complexes. Importantly, blockade of PIP_2_ microdomain formation significantly reduced the bulk endosome localization of Fwe (Figure 6d-f), although the SV localization of Fwe was also somewhat reduced (Figure 6d-e) Therefore, in addition to initiating ADBE, PIP_2_ microdomains can facilitate the retrieval of Fwe to the bulk endosome, enabling ADBE to remove its trigger by a negative feedback regulation and reducing endocytosis to prevent excess membrane uptake.

## Discussion

ADBE occurs immediately after exocytosis to retrieve required SV protein and lipid constituents to further regenerate SVs under conditions of high-frequency stimulations. Here, we show that the Fwe Ca^2+^ channel-dependent compartmentalization of PIP_2_ orchestrates coupling of exocytosis to ADBE and subsequent SV reformation. Based on our findings, we propose a model for this interplay (depicted in Figure 7). Under conditions of strong stimulation, SV exocytosis transfers Fwe from SVs to the periactive zone, where some of the activated Fwe provides the low Ca^2+^ levels that initiate Calcineurin activation to upregulate PIP_2_ (Step 1). Increased PIP_2_ enhances Fwe Ca^2+^ channel activity, thereby establishing a positive feedback loop that induces PIP_2_ microdomain formation (Step 2). High levels of PIP_2_ within these microdomains elicit bulk membrane invagination by triggering actin polymerization (Step 3). In parallel, PIP_2_ recruits adaptor protein complexes to facilitate proper sorting of Fwe to the bulk endosome (Step 4), thereby terminating the ADBE process. Finally, PIP_2_ microdomains dictate SV reformation from the bulk endosomes (Step 5), coordinating ADBE and subsequent SV reformation.

**Figure 7.**
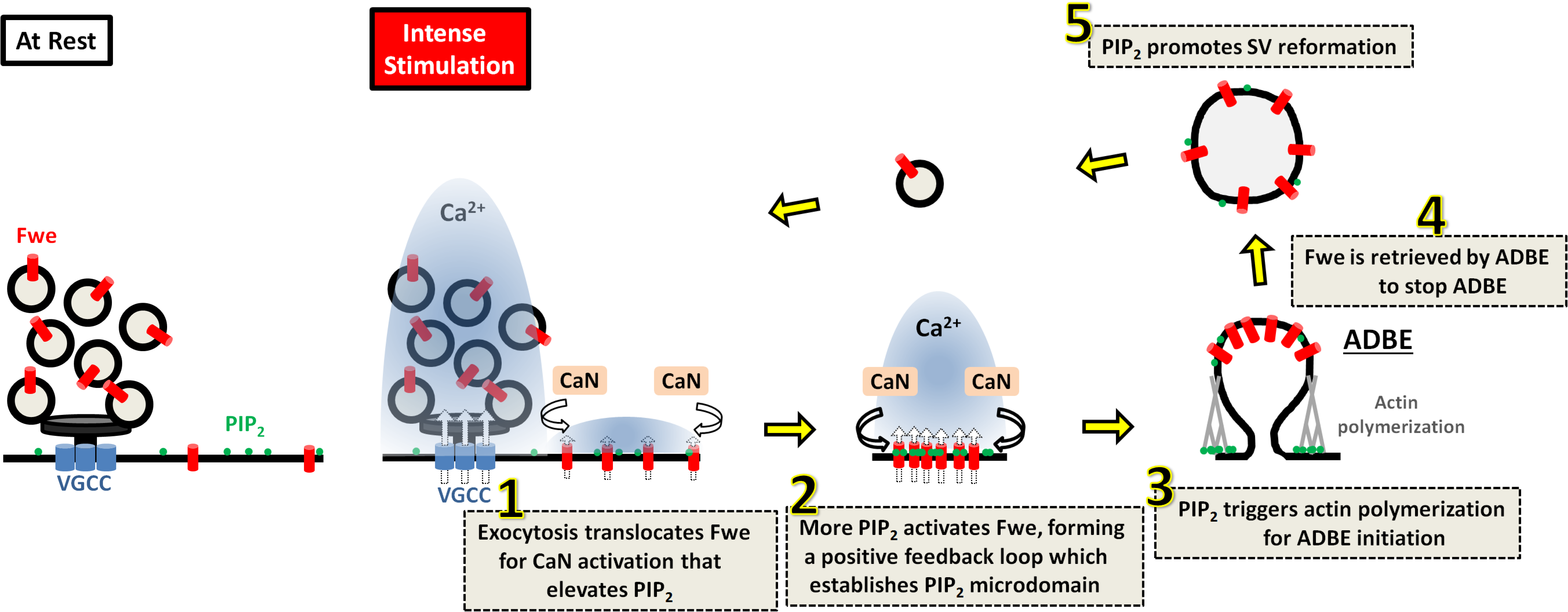
A proposed model for the role of Fwe-dependent PIP_2_ microdomains in coordinating ADBE and SV reformation from bulk endosomes. Details are described in the Discussion section.

### Fwe-dependent PIP_2_ microdomains trigger ADBE

The role of actin polymerization in ADBE has been investigated in mammals (Kononenko et al., 2014; Soykan et al., 2017; Wu et al., 2016), as well as in *Drosophila* (Akbergenova and Bykhovskaia, 2009). PIP_2_ is known to control a range of actin regulators, thereby modulating the dynamics of actin polymerization and branching (Janmey et al., 2018). It has been shown previously that, in response to nicotine stimulation, PIP_2_ forms microdomains of ~1 μm in diameter prior to the appearance of an actin-based ring structure in bovine chromaffin cells (Gormal et al., 2015). In agreement with this observation, we show that intense activity stimulation drives the formation of PIP_2_ microdomains at the periactive zone of *Drosophila* NMJ synaptic boutons. Perturbations of the formation of these microdomains reduces ADBE very significantly, demonstrating that rapid accumulation of PIP_2_ in microdomains is needed to trigger extensive actin polymerization, which likely generates sufficient mechanical force to produce the large endosomes. Furthermore, loss of *fwe* or *calcineurin* RNAi knockdown effectively inhibited the PIP_2_ microdomain formation and, as a consequence, ADBE. Those results are consistent with previous data arguing that Ca^2+^ promotes ADBE by activating its sensor Calcineurin (Cousin and Robinson, 2001; Jin et al., 2019a; Sun et al., 2010; Wu et al., 2009; Wu et al., 2014b; Xue et al., 2011).

Synj is a substrate of Calcineurin (Cousin and Robinson, 2001; Lee et al., 2004). Our data argue that the Fwe-derived Ca^2+^ activates Calcineurin to regulate PIP_2_ dynamics, most likely by modulating the functions of this PIP_2_-metabolizing enzyme. Under conditions of hyperosmotic stress, yeast cells undergo bulk membrane invagination, a process reminiscent of neuronal ADBE (Guiney et al., 2015). However, in that process, Calcineurin dephosphorylates Synj to alter its association with other endocytic partners rather than affecting its enzymatic activity to regulate PIP_2_ distribution. In addition, divalent Ca^2+^ ions are known to control lateral organization of PIP_2_, further compartmentalizing PIP_2_ into ~70 nm-sized microdomains in monolayers of lipid bilayers via electrostatic interactions between Ca^2+^ and PIP_2_ (Carvalho et al., 2008; Ellenbroek et al., 2011; Levental et al., 2009; Sarmento et al., 2014; Wang et al., 2012; Wen et al., 2018). Similarly-sized PIP_2_ clusters have been observed in PC12 cells under non-stimulated conditions (van den Bogaart et al., 2011). Therefore, temporal control of Calcineurin and Synj localization and Ca^2+^-mediated PIP_2_ clustering may account for the Fwe-dependent formation of PIP_2_ microdomains at the periactive zone prior to ADBE. Further investigations are needed to characterize the underlying mechanisms.

Our data show that direct binding of PIP_2_ is required for the Ca^2+^ channel activity of Fwe. Perturbation of PIP_2_-Fwe binding further impaired the formation of PIP_2_ microdomains as well as ADBE initiation. Hence, PIP_2_ controls Fwe gating, so that Fwe can promote PIP_2_ compartmentalization through a positive feedback regulation. Furthermore, loss of Fwe impaired intracellular Ca^2+^ increase evoked upon strong activity stimulation (Figure 3g) (Yao et al., 2017). These results argue that, in addition to PIP_2_, the channel function of Fwe may be gated by a significant change in membrane potential. In this notion, it is therefore possible that both factors may gate Fwe and thereby only allow channel opening when exocytosis locates Fwe to periactive zones. Future studies should explore the details of this channel gating mechanism.

### PIP_2_ microdomains coordinate retrieval of SV membranes and proteins for SV reformation via ADBE

Since ADBE is triggered very rapidly by intense stimuli, it was thought that this type of recycling randomly retrieves SV proteins and that the sorting process takes place when SVs regenerate from bulk endosomes. However, recent work has highlighted a distinct retrieval route for SV proteins during ADBE (Kokotos et al., 2018; Nicholson-Fish et al., 2015). Very little is known about the mechanisms underlying that retrieval route. Interestingly, removing VAMP4 or mutating its di-leucine motif was shown to impair ADBE (Nicholson-Fish et al., 2015). The di-leucine motif of transmembrane proteins is known to mediate binding with the AP-2 adaptor complex (Traub and Bonifacino, 2013). Given that the AP-2 adaptor complex works closely with PIP_2_ (McMahon and Boucrot, 2011), these findings imply a role for PIP_2_ and the AP-2 adaptor complex in SV protein sorting via ADBE. Indeed, our data show that bulk endosomes predominantly recycle Fwe, which depends on PIP_2_ microdomain-dependent recruitment of AP-2. Hence, in addition to initiating ADBE, PIP_2_ may participate in SV protein sorting to bulk endosomes.

SV regeneration occurs following formation of the bulk endosome. Our results also show that either removing Fwe-derived Ca^2+^ or perturbing PIP_2_ activity impaired the ability of SVs to reform from the bulk endosome, highlighting the essential role of PIP_2_ microdomains in this process. How could PIP_2_ of the plasma membrane regulate subsequent SV reformation? It has been shown that PIP_2_ is rapidly downregulated on bulk endosomes once formed by ADBE (Chang-Ileto et al., 2011; Cremona et al., 1999; Milosevic et al., 2011). It is conceivable that the high concentrations of PIP_2_ in microdomains may compensate for rapid turnover, thereby ensuring appropriate concentrations of PIP_2_ or PI(4)P for further recruitment of clathrin and adaptor protein complexes, such as AP-1 and AP-2 (Blumstein et al., 2001; Cheung and Cousin, 2012; Faundez et al., 1998; Glyvuk et al., 2010; Kokotos et al., 2018; Kononenko et al., 2014; Park et al., 2016). Alternatively, PIP_2_ may facilitate SV protein sorting prior to ADBE, meaning proper SV protein compositions on bulk endosomes could control recruitment of adaptor protein complexes. Both of these potential mechanisms are not mutually exclusive and may operate in parallel. Therefore, we propose that the Fwe-dependent formation of PIP_2_ microdomains is potentially important in coordinating retrieval of SV membranes and cargos when SVs are recycled via ADBE. Notably, the Fwe channel is evolutionarily conserved from yeast to human (Yao et al., 2009). We have also previously demonstrated conserved functions of Fwe in CME and ADBE at the mammalian central synapse (Yao et al., 2017). A recent study has also identified Fwe as a key protein mediating Ca^2+^-dependent granule endocytosis in mouse cytotoxic T lymphocytes (Chang et al., 2018). Hence, we hypothesize that the mechanism of ADBE we report here may be generally deployed across synapses and species, even in other non-neuronal cells.

## Materials and methods

### Drosophila strains and genetics

*fwe* mutants and transgenes *fwe^DB25^* and *fwe^DB56^* (Yao et al., 2009); *UAS-Flag-Fwe-HA* (Yao et al., 2009); *UAS-canA1FB5* (Dijkers and O’Farrell, 2007); *UAS-synj* (Khuong et al., 2013); *UAS-PLC_δ1_-PH-EGFP* (Verstreken et al., 2009)(Bloomington Drosophila Stock Center, BDSC#39693); *nSyb-GAL4* (Pauli et al., 2008); *elav-GAL4* (Bloomington Drosophila Stock Center, BDSC #8765); *vglut-lexA* (Bloomington Drosophila Stock Center, BDSC #60314); and *13XLexAop2-IVS-GCaMP6f* (Bloomington Drosophila Stock Center, BDSC #44277). Fly stocks were reared on regular food at 25 °C.

### Molecular cloning and transgenesis

For pUAST-HA-Fwe[WT]-APEX2, the coding sequence of the Fwe B isoform and APEX2 were separately PCR-amplified from pUAST-Flag-Fwe-HA (Yao et al., 2009) or pcDNA3 APEX2-NES (addgene #49386), respectively, and then subcloned into the pUAST vector. pUAST-HA-Fwe[K29A/R33A]-APEX2-HA was generated from pUAST-HA-Fwe[WT]-APEX2 through site-directed mutagenesis. pUAST-Flag-Fwe [K29/R33/K95/K100/R105/K146A/K147A/R150A]-HA, pUAST-Flag-Fwe[K146A/K147A/R150A]-HA and pUAST-Flag-Fwe[K95/K100/R105A]-HA were generated from pUAST-Flag-Fwe-HA through site-directed mutagenesis. For plexAOP2-PLC_δ1_-PH-APEX2-HA, the coding sequence of the PLC_δ1_-PH domain and APEX2-HA were separately PCR-amplified and then subcloned into pJFRC19-13XLexAop2-IVS-myr:GFP (addgene #26224). For YeMP-Nluc-Fwe-1D4, the cDNA fragment of Nluc and Fwe-1D4 were PCR-amplified from pNL1.1[Nluc] (a gift from Yi-Shiuan Huang) and YeMP-Fwe-1D4 plasmid (Yao et al., 2009), respectively, and then subcloned into the yeast expression YeMP vector. For YeMP-Nluc-Fwe[K29/R33A]-1D4, YeMP-Nluc-Fwe[K95/K100/R105/K146/K147/R150A]-1D4, and YeMP-Nluc-Fwe[K29/R33/K95/K100/R105/K146/K147/R150A]-1D4, the DNA fragments of the Fwe variant were PCR-amplified from above pUAST plasmids and sublconed to the YeMP-Nluc-Fwe-1D4 vector. Transgenic flies were made by Wellgenetics Inc.

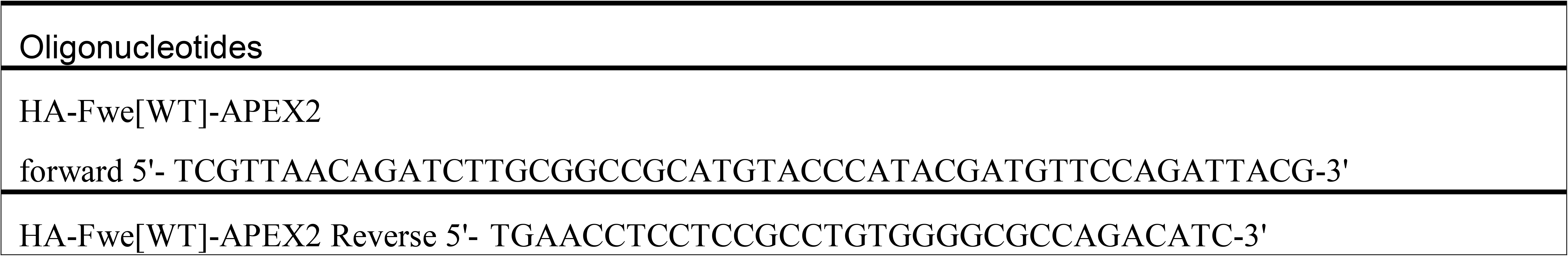
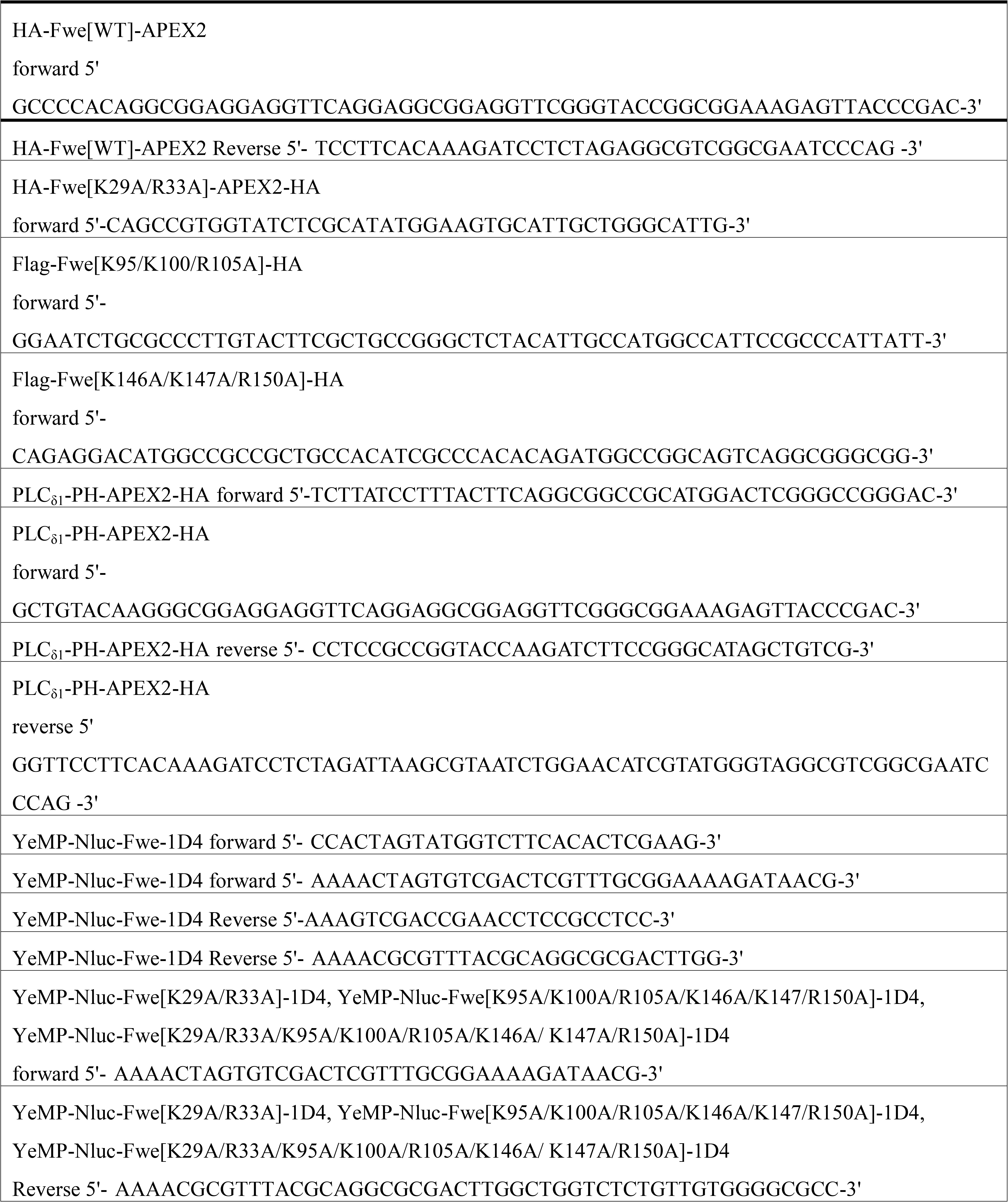

### Immunohistochemistry

Third instar larvae were fixed with 4% paraformaldehyde for 20 minutes. We used 1xPBS buffer containing 0.1% Tween-20 to stain the HA-tagged Fwe proteins. We used 1xPBS buffer containing 0.1 % Triton × 100 to stain PLC_δ1_-PH-EGFP or AP-2α. We used 1xPBS buffer containing 0.2% Triton × 100 to stain GCaMP6f. Primary antibodies were used as follows: chicken α-GFP (Invitrogen, 1:500); mouse α-HA (Sigma, 1:400), mouse α-Bruchpilot (Developmental Studies Hybridoma Bank nc82, 1:100); rabbit α-AP-2α (1:3000) (Gonzalez-Gaitan and Jackle, 1997); rabbit α-HRP conjugated with Alexa Fluor® 488, Cy3 or Cy5 (Jackson ImmunoResearch Laboratories, 1:250). Secondary antibodies conjugated to Alexa Fluor® 488, Alexa Fluor® 555, or Alexa Fluor® 647 (Invitrogen and Jackson ImmunoResearch) were used at 1:500. The NMJ boutons were derived from muscles 6 and 7 of abdominal segments 2/3. Images were captured with a Zeiss 780 confocal microscope and processed using LSM Zen and Image J software (National Institutes of Health).

### Biochemistry

For Western blotting, the brain and ventral nerve chord of larval fillets were removed and subjected to different stimulation conditions. Afterwards, the fillets were crushed in 1xSDS sample buffer and boiled for 10 minutes. For co-immunoprecipitation, liquid N_2_-frozen adult heads were crushed and homogenized in buffer K (150 mM NaCl, 10 mM HEPES (pH 7.4), 0.1 mM MgCl_2_, and 0.1 mM CaCl_2_) containing Complete Protease Inhibitor (Roche). Tissue debris was precipitated by centrifugation at 1000 × *g* for 1 min. The supernatant was solubilized in 1% Triton X-100 at 4 ºC for 1 h and subsequently centrifuged at 50,000 × *g* at 4 °C for 30 min in order to clean up the lysates. The resulting supernatant was incubated overnight with normal goat serum-adsorbed IgG beads (Sigma) and then with α-HA-coupled beads (Sigma) or RFP TrapA beads (Chromotek). The beads were washed with buffer (100 mM NaCl, 10 mM HEPES pH 7.4) and boiled in 1xSDS sample buffer for 10 min. Dilutions for primary antibodies were as follows: rabbit anti-AP-2α, 1:10000 (Gonzalez-Gaitan and Jackle, 1997); mouse anti-α-actin, 1:50000 (Sigma); chicken anti-GFP, 1:5000 (Invitrogen).

### PLA

Third instar larvae were fixed with 4% paraformaldehyde for 20 min and permeabilized with 1xPBS buffer containing 0.1% Tween-20. Larval fillets were incubated with mouse α-HA (Sigma, 1:200) and rabbit α-GFP (Invitrogen, 1:500) in 1xPBS buffer containing 0.1% Tween-20 at 4 °C for 12 h. Excess antibodies were washed out using 1xPBS buffer containing 0.1% Tween-20. The samples were mixed with the PLA probe (Sigma, 1:5) for 2 h at 37 °C. After washing with 1xbuffer A, the samples were incubated with ligation solution (1:40) for 1.5 h at 37 °C. After washing with 1xbuffer A, the samples were incubated with amplification solution (1:80) for 2 h at 37 °C. Next, the samples were washed with 1xbuffer B and then 0.01xbuffer B. The samples were stained with anti-chicken Alexa Fluor® 488-conjugated IgG and anti-mouse Alexa Fluor® 647-conjugated IgG, followed by a wash of 1xPBS buffer containing 0.1% Tween-20. The samples were then mounted for confocal imaging.

### Nluc-Fwe-1D4 purification and BRET assay

The Nluc-Fwe-1D4 fusion protein was purified as described previously (Yao et al., 2009). Briefly, plasma membrane was isolated from a 2 liter culture of yeast strain BJ5457 expressing Nluc-Fwe-1D4 protein. The plasma membranes were solubilized at 4 °C with 10x critical micelle concentration (CMC) DDM (Anatrace) in a solution of 20mM HEPES (pH8.0), 300 mM NaCl, 10% Glycerol, 2.0 mM DTT, 1mM PMSF for 2 h. Insoluble membranes were spun down by centrifugation at 100,000 × *g* for 60 min. The lysates including solubilized Nluc-Fwe-1D4 proteins were cleaned with CNBr sepharose 4B at 4 °C for 1 h. The samples were then mixed with α-1D4-conjugated CNBr-activated Sepharose 4B at 4 °C for 8-12 h. After washing with a solution of 20mM HEPES (pH8.0), 150 mM NaCl, 10% Glycerol, 2.0 mM DTT, 1mM PMSF and 2.6xCMC DDM, the protein was eluted with a 1D4 peptide-containing buffer (3 mg of 1D4 peptide in 1 ml of washing buffer). Purified proteins were subjected to SDS-PAGE and detected using Lumitein staining or Western blotting with anti-α-1D4 antibody at 1:5000 (Yao et al., 2009). The BRET assay was performed in a 384-well plate, with each well containing 30 μl of reaction solution [0.5 nM purified proteins, 5 μM BODIPY-TMR Phosphatidylinositol 4,5-bisphosphate (C-45M16A, Echelon Bioscience), furimazine (Promega, 1:2000), 20 mM HEPES, 150 mM NaCl, 10% Glycerol, 2 mM DTT, 1 mM PMSF, 4 mM DDM]. Fluorescence signal was detected using a Microplate Reader M1000 pro (Tecan) with two different emission spectrum filters, i.e., 500-540 nm for Nluc and 550-630 nm for BODIPY-TMR Phosphatidylinositol 4,5-bisphosphate. For competition assay, 30 μl of the reaction solution was included with 1 mM brain phosphatidylinositol 4,5-bisphosphate (Avanti). The BRET signal was calculated according to the following formula:

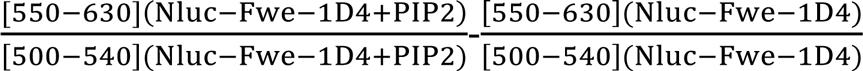

### Live imaging

To visualize PLC_δ1_-PH-EGFP signal, third instar larvae were dissected in a zero-calcium HL-3 solution at room temperature. For groups stimulated with electric pulses, larval fillets were bathed in a solution of 2 mM Ca^2+^ (70 mM NaCl, 5 mM KCl, 10 mM MgCl_2_, 10 mM NaHCO_3_, 5 mM trehalose, 5 mM HEPES (pH 7.4), 115 mM sucrose, 2 mM CaCl_2_). High concentrations of glutamate were used to desensitize glutamate receptors, thereby reducing muscle contraction when stimulated. A cut axonal bundle was sucked into the tip of a glass capillary electrode and then stimulated at 20 or 40 Hz for 3 min. Stimulus strength was set at 5 V and 0.5 ms duration by means of pClamp 10.6 software (Axon Instruments Inc.). After stimulation for two min, muscle contraction significantly decelerated. Thus, we captured individual snapshot images every second from the third minute. Muscles 6 and 7 of abdominal segment 3 were imaged. Under the condition of high K^+^ stimulation, larval fillets were bathed in a solution of 90 mM K^+^/2 mM Ca^2+^/7 mM glutamate (25 mM NaCl, 90 mM KCl, 10 mM MgCl_2_, 10 mM NaHCO_3_, 5 mM trehalose, 5 mM HEPES (pH 7.4), 30 mM sucrose, 2 mM CaCl_2_, 7 mM monosodium glutamate) for 5 min. Images were taken from the fifth minute of stimulation to the end of stimulation. Imaging was done using MetaMorph software and an ANDOR iXon3 897 camera. To visualize GCaMP6f signal, third instar larvae were dissected in a zero-calcium HL-3 solution at room temperature. Images were taken of larval fillets at rest. Subsequently, larval fillets were stimulated with a solution of 90 mM K^+^/2 mM Ca^2+^/7 mM glutamate for 5 min. Imaging was captured from the fifth minute of stimulation to the end of stimulation, with one image captured per second. We used the *lexA/lexAOP2* binary system to stably express a comparable level of GCaMP6f in the presynaptic compartment of the NMJ bouton among the genotypes tested, allowing us to compare Ca^2+^ imaging results when the resting Ca^2+^ levels were potentially affected by genetic backgrounds. We did immunostaining of the GCaMP6f protein with a-GFP antibody. A comparable GCaMP6f level among different genotypes tested in each dataset was confirmed. Evoked Ca^2+^ levels were calculated by subtracting the resting GCaMP6f fluorescence from the GCaMP6f fluorescence induced by high K^+^ stimulation. The NMJs were derived from muscles 6 and 7 of abdominal segment 2/3. Image processing and quantification were achieved using LSM Zen and Image J.

### FM dye assay

For the FM4-64 dye loading/unloading assay, third instar larvae were dissected in zero Ca^2+^ HL-3 solution at room temperature and stimulated with a HL-3 solution of 90 mM K^+^/2 mM Ca^2+^/4 μM FM4-64 (Invitrogen) for 5 min to load FM4-64 dye into the boutons. Excess dye was removed by extensive washing of zero-calcium HL-3 solution. FM1-43 dye uptake of the boutons was imaged and quantified to indicate “Loading”. The loaded dye in SVs was unloaded via 1-min stimulation of 90 mM K^+^/2 mM Ca^2+^. Released dye was washed out with zero-calcium HL-3 solution. The residual dye represented “Unloading”. The efficacy of exocytosis was calculated as (F_Loading_-F_Unloading_)/F_Loading_. Images were captured with a Zeiss 780 confocal microscope. To quantify clathrin-mediated endocytosis, third instar larvae were dissected and stimulated with a HL-3 solution of 90 mM K^+^/0.5 mM Ca^2+^/4 μM fixable FM1-43 (Invitrogen) for 1 min, followed by extensive washing of zero-calcium HL3 solution for 10 min and fixation for 10 min. Images were captured with a Zeiss 780 confocal microscope. Image processing and quantification were achieved using LSM Zen and Image J.

### Transmission electron microscopy

Third instar larval fillets were prepared in zero-calcium HL-3 solution at room temperature. For the resting conditions, the fillets were bathed in zero-calcium HL-3 solution at room temperature for another 10 min before fixation. For the high K^+^ stimulation conditions, fillets were bathed in a solution of 90 mM K^+^ and 2 mM Ca^2+^ for 10 min. The stimulation was terminated by washing three times with zero-calcium HL-3 solution, followed by fixation. For recovery conditions, following high K^+^ stimulation, fillets were bathed in zero-calcium HL-3 solution at room temperature for 10 or 20 min before fixation. Larval fillets were fixed for 12 h at 4 °C in 4% paraformaldehyde/1% glutaraldehyde/0.1 M cacodylic acid (pH 7.2), rinsed with 0.1 M cacodylic acid (pH 7.2), and postfixed with 1% OsO_4_ and 0.1 M cacodylic acid at room temperature for 3 h. For diaminobenzidine (DAB) polymerization, third instar larvae were dissected at room temperature in zero-calcium HL-3 medium, followed by a 10-min incubation in 5 mM K^+^/0 Ca^2+^ mM solution or a 10-min stimulation of 90 mM K^+^/2 mM Ca^2+^. Next, the samples were subjected to 30-min fixation in ice-cold 4% paraformaldehyde/1% glutaraldehyde/0.1 M cacodylic acid (pH 7.2). Subsequently, the samples were transferred to eppendorf tubes for 15-min incubation with a solution of 0.5 mg/ml DAB solution, followed by incubation with a solution of 0.5 mg/ml DAB and 0.006% H_2_O_2_ for 15 min at room temperature. This latter step was repeated once to ensure DAB polymerization. Samples were washed three times with 1xPBS buffer for 10 min and then fixed with a solution of 4% paraformaldehyde/1% glutaraldehyde/0.1 M cacodylic acid (pH 7.2) for 12 h at 4 ºC, followed by fixation with a solution of 1% OsO_4_/0.1 M cacodylic acid at room temperature for 3 h. These samples were then subjected to a series of dehydration steps using 30-100% ethanol. After 100% ethanol dehydration, the samples were sequentially incubated with propylene, a mixture of propylene and resin, and pure resin. Finally, the samples were embedded in 100% resin. TEM images were captured using Tecnai G2 Spirit TWIN (FEI Company) and a Gatan CCD Camera (794.10.BP2MultiScanTM). NMJ boutons were captured at high magnifications. Statistical analyses were performed using Image J.

### Statistics

All data analyses were conducted using GraphPad Prism 8.0, unless stated otherwise. Paired and multiple datasets were compared by Student *t*-test or one-way ANOVA with Tukey’s post hoc test, respectively. Individual data values are biological replicates. Samples were randomized during preparation, imaging, and data processing to minimize bias.

## Acknowledgments

We thank Hugo Bellen, Patrik Verstreken, Kartik Venkatachalam, the Bloomington Drosophila Stock Center, and the Developmental Studies Hybridoma Bank for stocks and reagents. We thank Hugo Bellen, Ruey-Hwa Chen, and Y. Henry Sun for the critical comments. We thank Yi-Shuian Huang for providing the Nluc plasmid. We thank Wellgenetics for making transgenic lines. We thank the IMB imaging core for helping with TEM and SIM imaging. This work was supported by grants from the Ministry of Science and Technology of Taiwan (107-2311-B-001-003-MY3, 106-0210-01-15-02, and 107-0210-01-19-01).

---

**Figure 1-figure supplement 1.**
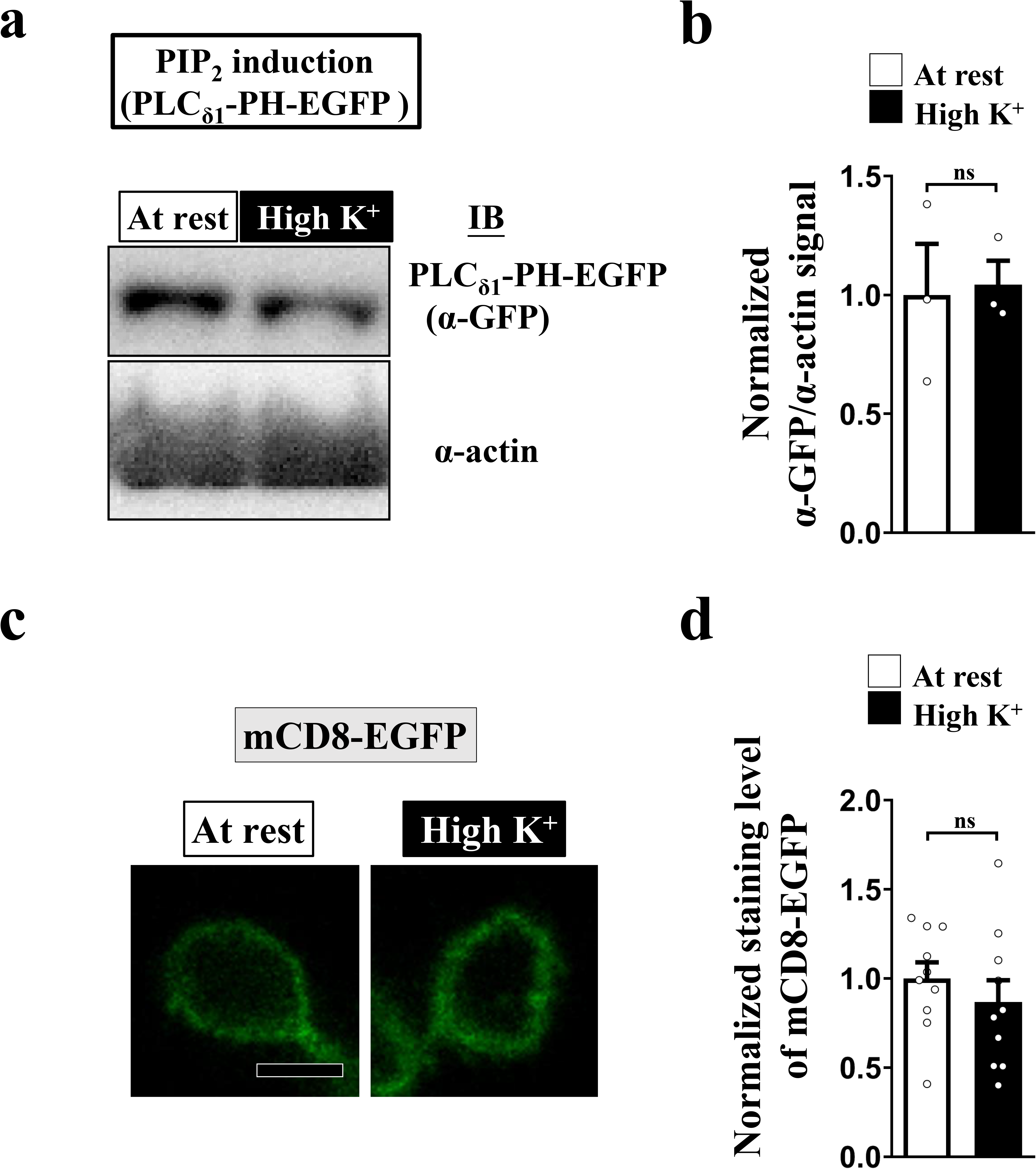
The change in the PLC_δ1_-PH-EGFP fluorescence is responsible for an increase in the PIP_2_ level. (a-b) The protein expression level of PLC_δ1_-PH-EGFP was not changed upon high K^+^ stimulation. (a) Larval fillets expressing *UAS-PLC_δ1_-PH-EGFP* using *nSyb-GAL4* were dissected, and the brain and ventral nerve cord were removed. The samples were subject to the resting condition (10-min incubation of 5 mM K^+^/0 mM Ca^2+^) or high K^+^ stimulation (10-min stimulation of 90 mM K^+^/2 mM Ca^2+^), and then immediately lysed in 1xSDS sample buffer. The lysate of a single larva was loaded in each lane of the Western blot, and α-GFP and α-actin were used to detect presynaptic levels of PLC_δ1_-PH-EGFP or to serve as loading control, respectively. Quantification data for the ratio of α-GFP to α-actin blotting signal, normalized to the value of the resting condition. (c) Confocal images of NMJ boutons expressing *UAS-mCD8-GFP* using *nSyb-GAL4*. The boutons were subjected to the resting condition (10-min incubation of 5 mM K^+^/0 mM Ca^2+^) or high K^+^ stimulation (10-min stimulation of 90 mM K^+^/2 mM Ca^2+^), followed by chemical fixation and α-GFP immunostaining. (d) Quantification data for mCD8-EGFP staining intensities, normalized to the value of the resting condition. Individual data values are shown in graphs. *P* values: ns, not significant. Mean ± SEM. Scale bar: 2 μm (c). Statistics: Student *t*-test (b, d). **Figure 1-figure supplement 1-source data 1. Source data for Figure 1-figure supplement 1**.

**Figure 2-figure supplement 1.**
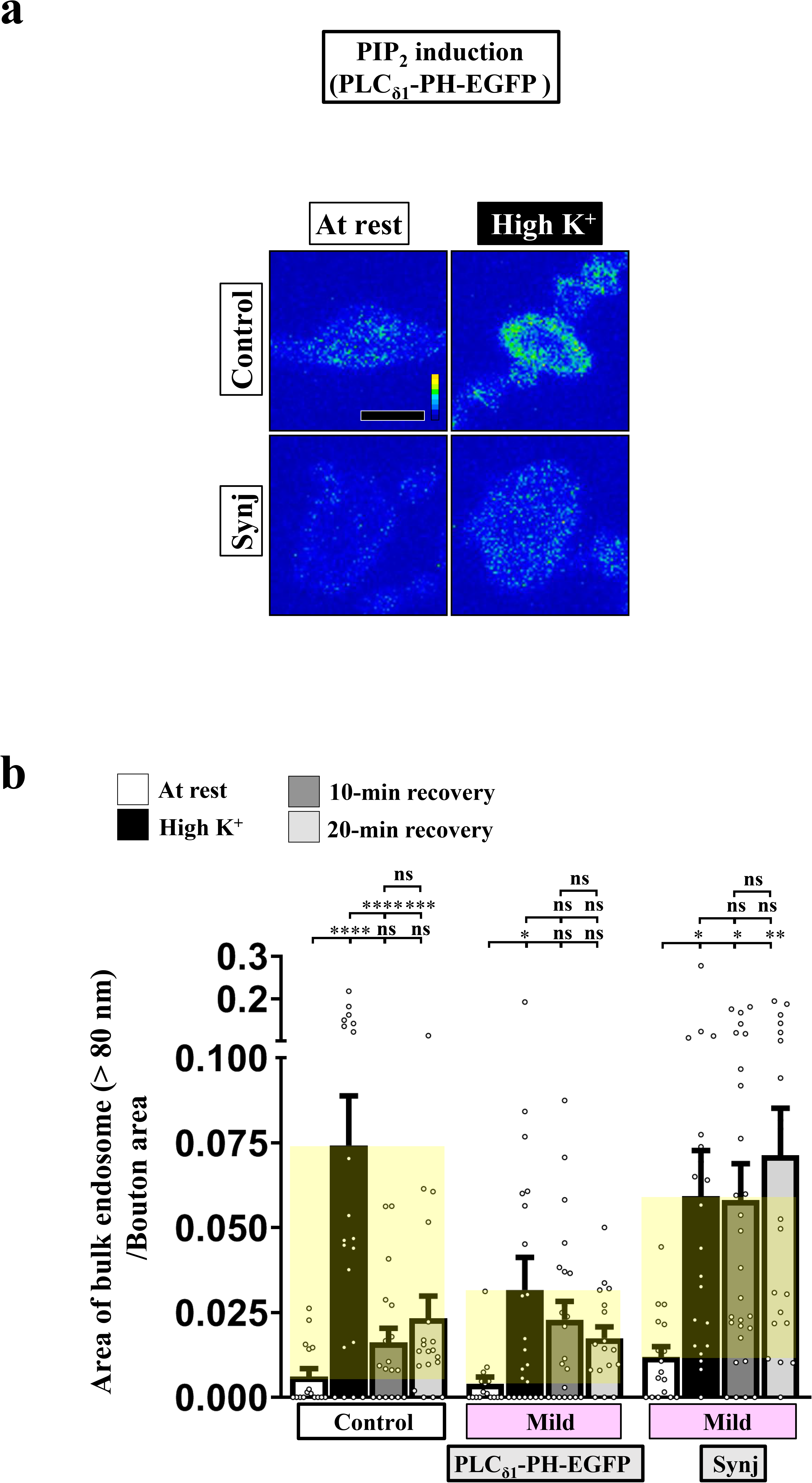
Blockade of the PIP_2_ microdomains suppresses ADBE. (a) Pseudocolored confocal images of NMJ boutons co-expressing *UAS-PLC_δ1_-PH-EGFP* with *UAS-RFP* control or *UAS-Synj*. The boutons were subjected to the resting condition (10-min incubation of 5 mM K^+^/0 mM Ca^2+^) or high K^+^ stimulation (10-min stimulation of 90 mM K^+^/2 mM Ca^2+^), followed by chemical fixation and α-GFP immunostaining. Scale bar: 2 μm. (b) Quantification data for the ratio of total area of bulk endosomes to bouton area. Indicated genotypes were assessed. Representative TEM images are shown in Figure 2a. Individual data values are shown in graphs. *P* values: ns, not significant; *, *P*<0.05; **, *P*<0.01; ***, *P*<0.001; ****, *P*<0.0001. Mean ± SEM. Statistics: one-way ANOVA with Tukey’s post hoc test. **Figure 2-figure supplement 1-source data 1. Source data for Figure 2-figure supplement 1**.

**Figure 2-figure supplement 2.**
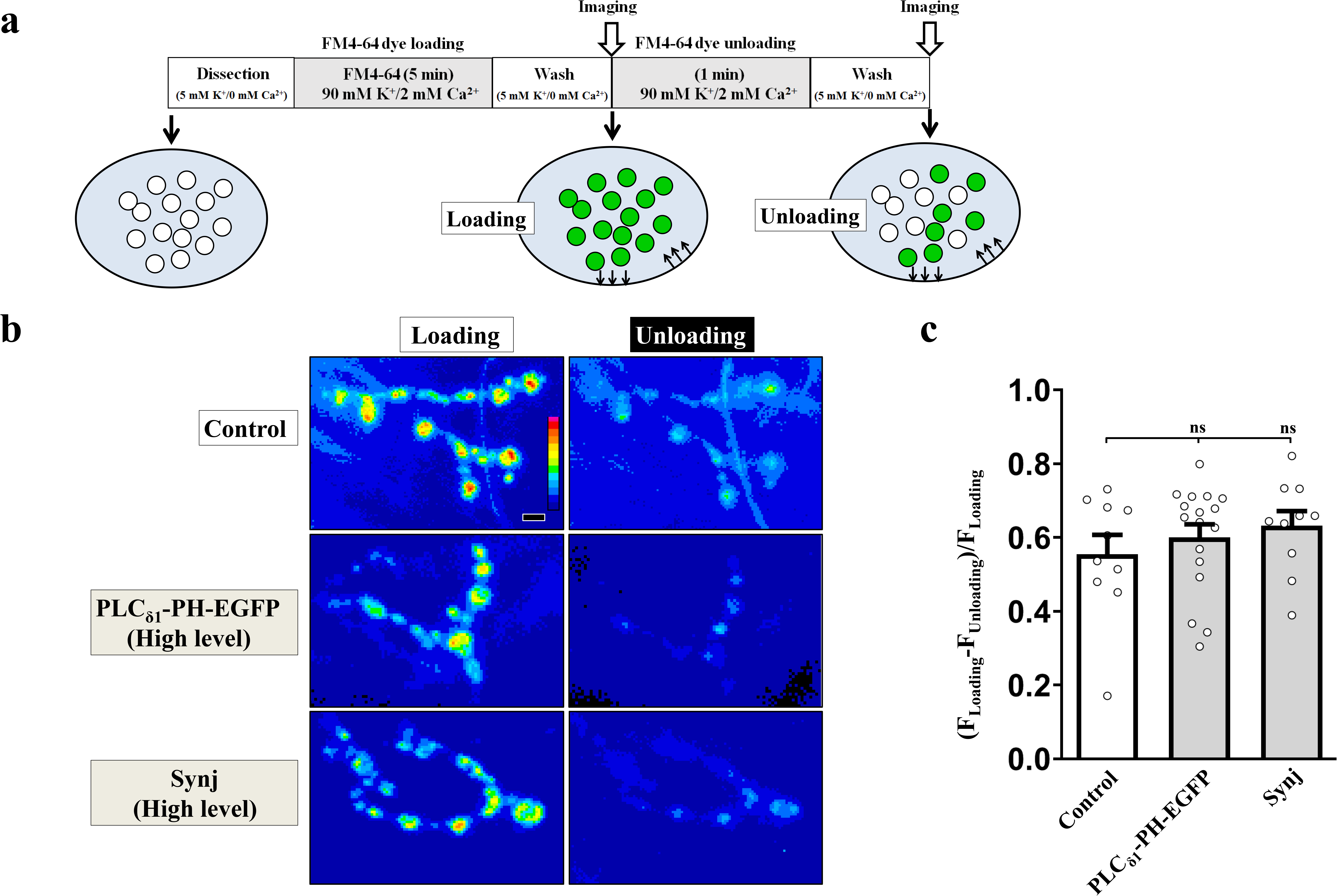
SV exocytosis, as measured by a FM4-64 loading/unloading assay. (a) Schematic of our FM4-64 dye loading/unloading assay. Larval fillets were prepared in a solution of 5 mM K^+^/0 mM Ca^2+^ and then subjected to 5-min stimulation of 90 mM K^+^/2 mM Ca^2+^ in the presence of 4 μM fixable FM4-64 dye. After extensive washing with a solution of 5 mM K^+^/0 mM Ca^2+^, the boutons were imaged to indicate dye loading of the SV. Dye unloading was performed by 1-min stimulation of 90 mM K^+^/2 mM Ca^2+^. After washing away released dye, the boutons were imaged to indicate dye unloading. (b) Pseudocolored images of FM4-64 dye-labeled boutons derived from control (*nSyb-GAL4*), high expression of PLC_δ1_-PH-EGFP (*nSyb-GAL4/UAS-PLC_δ1_-PH-EGFP* at 29 ºC), and high expression of Synj (*nSyb-GAL4/UAS-synj* at 29 ºC). (c) (F_Loading_-F_Unloading_)/F_Loading_ was calculated to indicate the efficacy of SV exocytosis. Individual data values are shown in the graphs. *P* values: ns, not significant. Mean ± SEM. Scale bar: 2 μm. Statistics: one-way ANOVA with Tukey’s post hoc test. **Figure 2-figure supplement 2-source data 1. Source data for Figure 2-figure supplement 2**.

**Figure 4-figure supplement 1.**
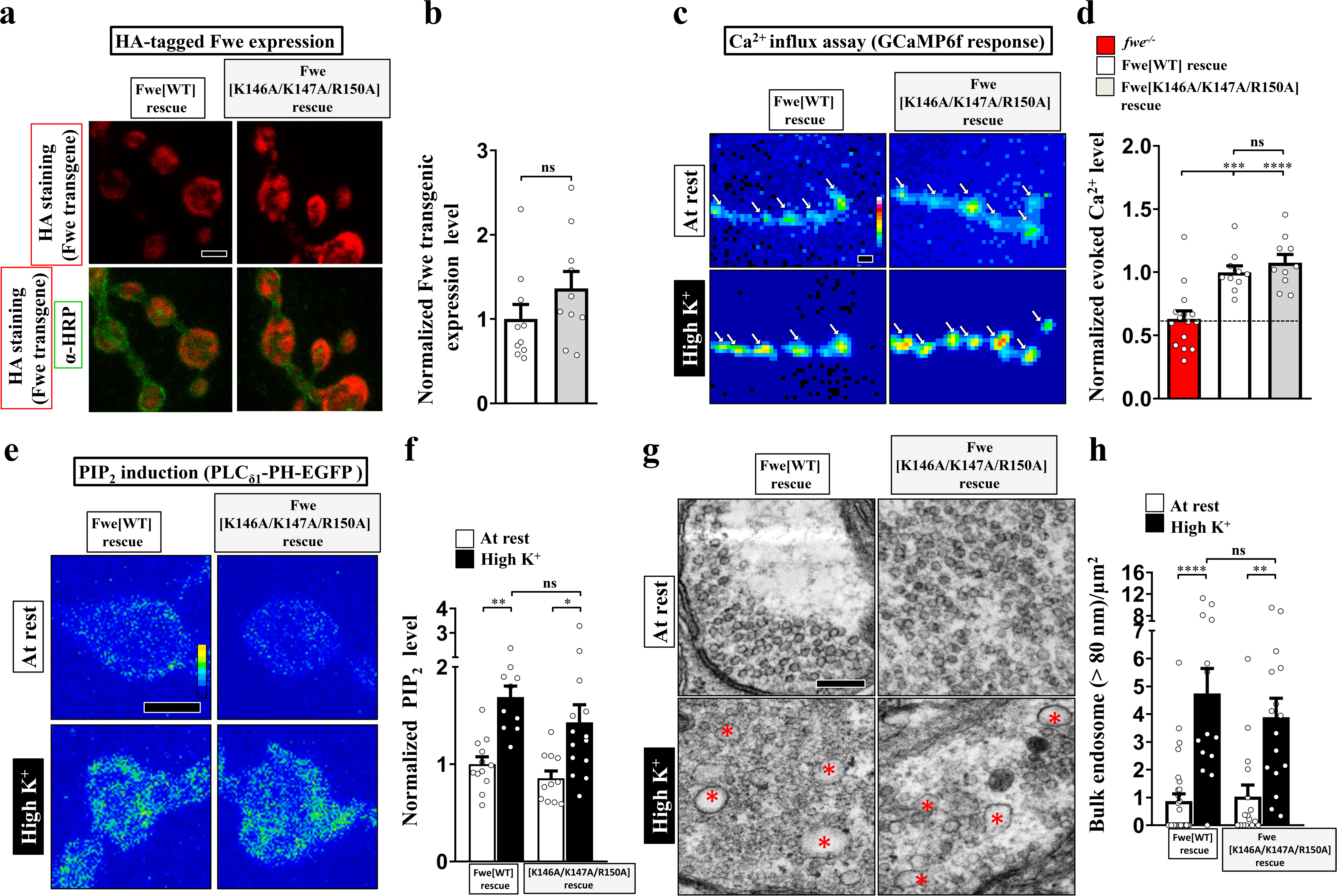
The C-terminal residues (K146/K147/R150) of Fwe are not involved in regulating Ca^2+^ channel activity. (a-b) Alanine substitution of the C-terminal residues K146/K147/R150 does not influence the expression or SV localization of Fwe. (a) Confocal images of NMJ boutons derived from the larvae of Flag-Fwe[WT]-HA rescue (*nSyb-GAL4/UAS-Flag-Fwe-HA in fwe^DB25/DB56^*) or Flag-Fwe[K146A/K147A/R150A]-HA rescue *(nSyb-GAL4/UAS-Flag-Fwe[K146A/K147A/ R150A]-HA in fwe^DB25/DB56^*) lines. α-HA (red) and α-HRP (green) staining was used to detect Fwe transgenic proteins and neuronal membranes, respectively. α-HRP staining was also used as the staining control. (b) Quantification data for the staining intensity ratio of α-HA to α-HRP, normalized to the value of the Flag-Fwe[WT]-HA rescue line. (c-h) Alanine substitution of the C-terminal residues K146/K147/R150 does not influence Ca^2+^ channel activity, the ability to induce PIP_2_ microdomains and ADBE. (c) Snapshot Ca^2+^ images of the NMJ boutons. *fwe^−/−^* (*fwe^DB25/DB56^*). The boutons were subjected to 5-min stimulation of 90 mM K^+^/2 mM Ca^2+^. The images were taken in the fifth minute. Evoked Ca^2+^ levels are represented as the increase in GCaMP6f fluorescence in response to high K^+^ stimulation. (d) Quantification data for evoked Ca^2+^ level, normalized with the value of the control. (e) Pseudocolored confocal images of the NMJ boutons. The boutons were subjected to the resting condition (10-min incubation of 5 mM K^+^/0 mM Ca^2+^) or high K^+^ stimulation (10-min stimulation of 90 mM K^+^/2 mM Ca^2+^), followed by chemical fixation and α-GFP immunostaining. (f) Quantification data for the PLC_δ1_-PH-EGFP staining intensity, normalized to the value of the resting condition of the Flag-Fwe[WT]-HA rescue line. (g) TEM images of NMJ boutons. TEM processing was performed under the following conditions: at rest (10-min incubation of 5 mM K^+^/0 mM Ca^2+^); or high K^+^ (10-min stimulation of 90 mM K^+^/2 mM Ca^2+^). Bulk endosomes (red asterisks). Mitochondria (mt). (h) Quantification data of total numbers of bulk endosomes per bouton area. Individual data values are shown in the graphs. *P* values: ns, not significant; *, *P*<0.05; **, *P*<0.01; ***, *P*<0.001; ****, *P*<0.0001. Mean ± SEM. Scale bar: 2 μm (a, c, e), 500nm (g). Statistics: Student *t*-test (b). One-way ANOVA with Tukey’s post hoc test (d, f, h). **Figure 4-figure supplement 1-source data 1. Source data for Figure 4-figure supplement 1**.

**Figure 4-figure supplement 2.**
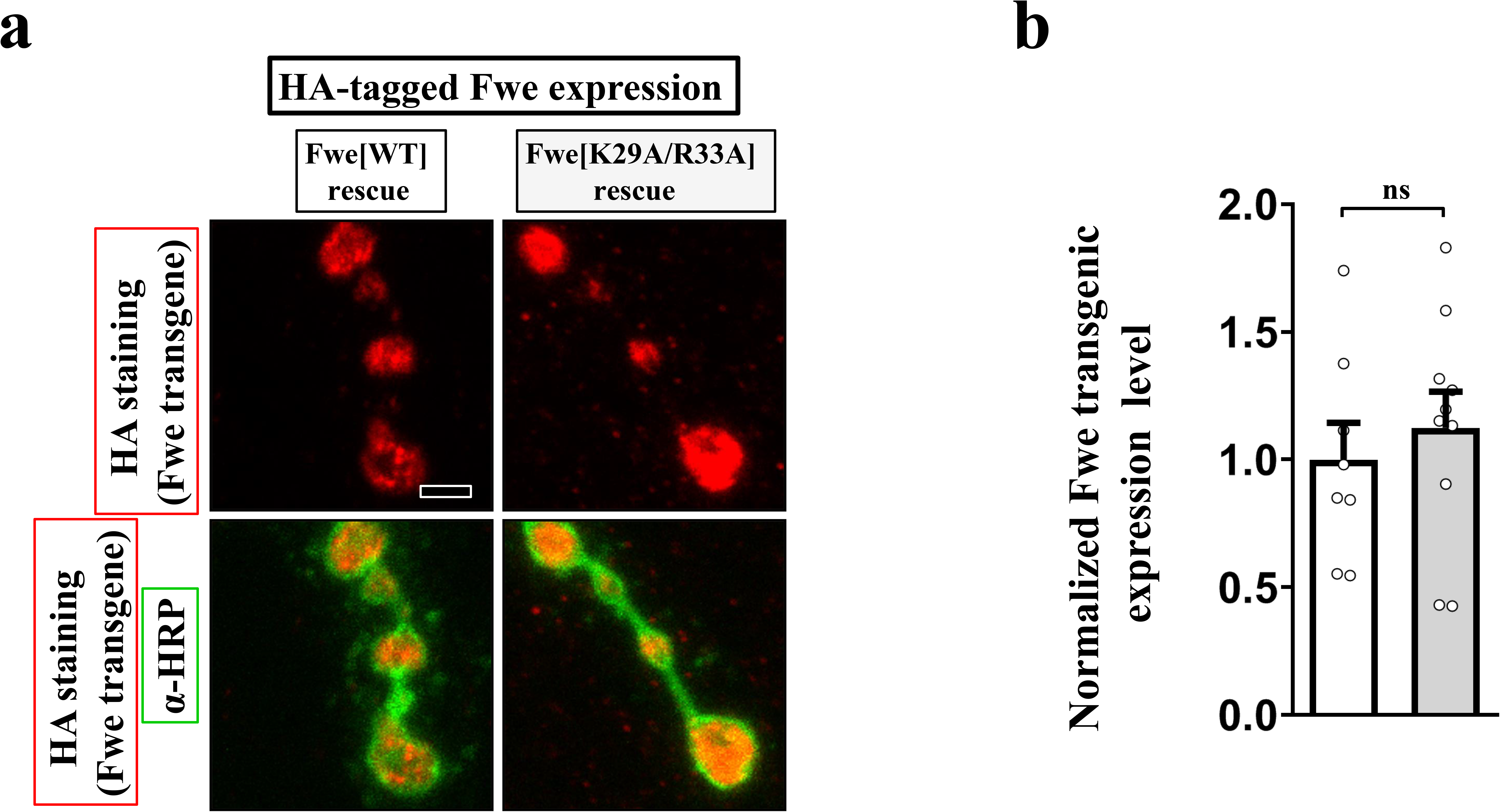
Expression of Fwe[R29A/K33A]. (a-b) The Fwe[K29/R33A] mutant variant properly localizes to the presynaptic compartment. (a) Confocal images of NMJ boutons derived from the larvae of the HA-Fwe[WT]-APEX2 rescue (*nSyb-GAL4/UAS-HA-Fwe[WT]-APEX2 in fwe^DB25/DB56^*) or HA-Fwe[K29/R33A]-APEX2 rescue *(nSyb-GAL4/UAS-HA-Fwe[K29/R33A]-APEX2 in fwe^DB25/DB56^*) lines. α-HA (red) and α-HRP (green) staining was used to detect Fwe transgenic proteins and neuronal membranes, respectively. α-HRP staining was also used as the staining control. (b) Quantification data for the staining intensity ratio of α-HA to α-HRP, normalized to the value of the Flag-Fwe[WT]-HA rescue line. Individual data values are shown in the graphs. *P* values: ns, not significant. Scale bar: 2 μm. Statistics: Student *t*-test. **Figure 4-figure supplement 2-source data 1. Source data for Figure 4-figure supplement 2**.

**Figure 5-figure supplement 1.**
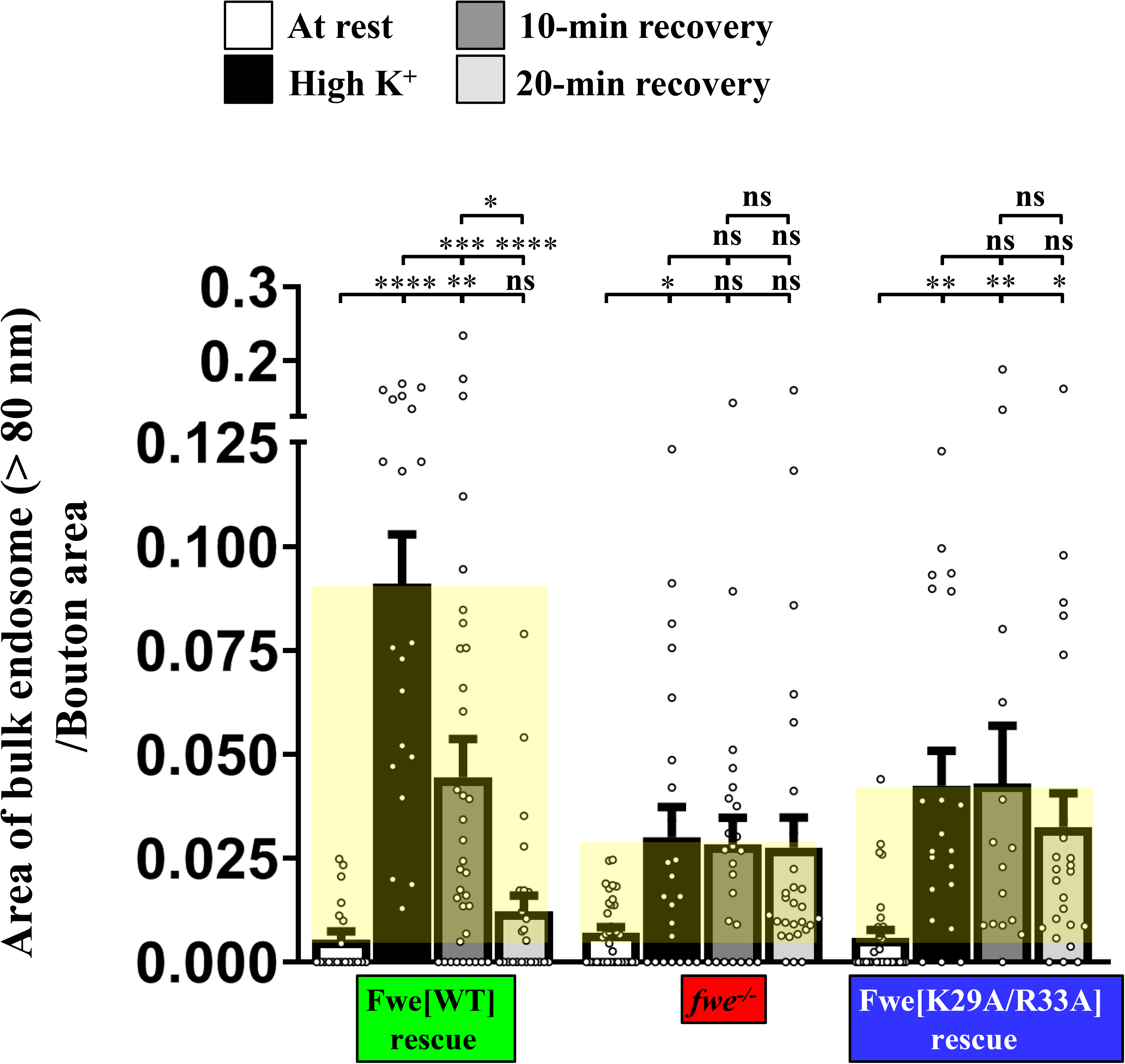
Blockade of the positive feedback loop between Fwe and PIP_2_ abolishes SV reformation from the bulk endosome. Quantification data for the ratio of total area of bulk endosomes to bouton area of the indicated genotypes. Individual data values are shown in the graphs. *P* values: ns, not significant; *, *P*<0.05; **, *P*<0.01; ***, *P*<0.001; ****, *P*<0.0001. Mean ± SEM. Scale bar: 2 μm. Statistics: One-way ANOVA with Tukey’s post hoc test. **Figure 5-figure supplement 1-source data 1. Source data for Figure 5-figure supplement 1**.

